# A metabolic pathway for glucosinolate activation by the human gut symbiont *Bacteroides thetaiotaomicron*

**DOI:** 10.1101/626366

**Authors:** Catherine S. Liou, Shannon J. Sirk, Camil A.C. Diaz, Andrew P. Klein, Curt R. Fischer, Steven K. Higginbottom, Justin L. Sonnenburg, Elizabeth S. Sattely

**Author notes:** These authors contributed equally (Catherine S. Liou, Shannon J. Sirk, Camil A.C. Diaz).

## Abstract

Diet is the largest source of plant-derived metabolites that influence human health. The gut microbiota can metabolize these molecules, altering their biological function. However, little is known about the gut bacterial pathways that process plant-derived molecules. Glucosinolates are well-known metabolites in brassica vegetables and metabolic precursors to cancer-preventive isothiocyanates. Here, we identify a genetic and biochemical basis for isothiocyanate formation by *Bacteroides thetaiotaomicron,* a prominent gut commensal species. Using a genome-wide transposon insertion screen, we identified an operon required for glucosinolate metabolism in *B. thetaiotaomicron.* Expression of BT2159-BT2156 in a non-metabolizing relative, *Bacteroides fragilis*, resulted in gain of glucosinolate metabolism. We show that isothiocyanate formation requires the action of BT2158 and either BT2156 or BT2157 *in vitro*. Monocolonization of mice with mutant *Bt*Δ*2157* showed reduced isothiocyanate production in the gastrointestinal tract. These data provide insight into the mechanisms by which a common gut bacterium processes an important dietary nutrient.

## INTRODUCTION

In addition to contributing fiber and essential micronutrients to our diet, plants are a rich source of secondary metabolites capable of eliciting pharmacological effects (Holst and Williamson, 2008; Martin et al., 2013). The vast diversity of these compounds in dietary plants has exceeded our understanding of both their metabolic fate following consumption and resulting effect on human physiology, limiting our ability to assess and optimize their effects on health. Commensal microbes in the gut have been implicated in the processing and activation of many of these plant-derived bioactive molecules, altering their chemical structures and reactivity, biochemical function, and propensity to be absorbed through the intestinal epithelium. However, these activities are often attributed to taxonomic groups and are rarely understood at the genetic or biochemical level (Bode et al., 2013; Bokkenheuser et al., 1987; Clavel et al., 2006). A detailed and mechanistic understanding of how gut microbial metabolism impacts host biology will yield insight into how, from a common dietary input, the microbiome can drive divergent physiological outcomes based on which nutrient processing pathways predominate.

A lead example of diet-derived small molecules that influence the host involves edible cruciferous vegetables such as broccoli, kale, cabbage, and many other plants in the family Brassicaceae. Epidemiological evidence has linked diets rich in cruciferous vegetables to a decreased risk of gastrointestinal cancer, and isothiocyanates (ITC) are thought to be responsible for the chemopreventive properties of these plants (Herr and Büchler, 2010). ITC do not typically accumulate in dietary crucifers; instead, they are hydrolysis products of biologically inert glucosinolates (GS), a class of thioglucosides with a sulfonated oxime moiety and an amino-derived side chain that are abundant in the plant tissue (Figure 1A). Recent efforts to take advantage of these protective effects have included the development and commercialization of Beneforte broccoli, a line bred to accumulate enhanced levels of the methionine-derived GS glucoraphanin (Mithen, 2013).

**Figure 1.**
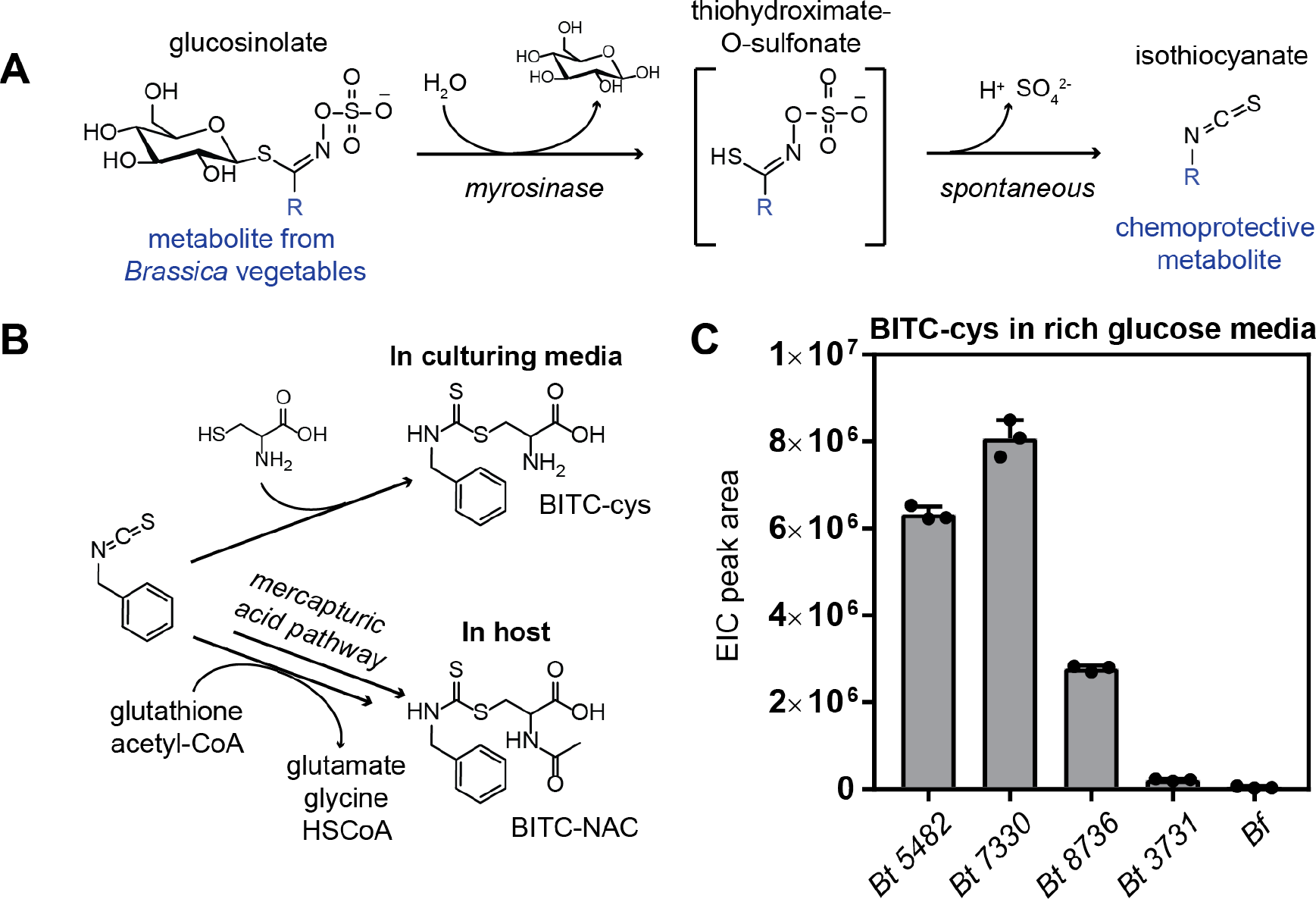
Activation of glucosinolates to isothiocyanates by microbial myrosinases. (A) Reaction scheme for the conversion of glucosinolates to isothiocyanates by myrosinases (B) Metabolic fates of microbially produced isothiocyanates (benzyl isothiocyanate, derived from the glucotropaeolin, shown) in the culturing media or in the host urine. Isothiocyanate in culturing media was measured as ITC-cysteine, derived from *in situ* conjugation with cysteine in the media. Isothiocyanate in host urine was measured as the N-acetyl cysteine conjugate, an excreted product of mercapturic acid pathway metabolism of glutathione-conjugated isothiocyanate (Hwang and Jeffery, 2003). **(C)**Benzyl isothiocyanate cysteine conjugate produced from glucotropaeolin by different *Bacteroides* strains cultured in rich glucose media for 24 hours. Data shown is the mean ± SD of three biological replicates. An LC-MS extracted ion chromatogram (*m/z* 271.0570) was used to track quantities of BITC-cys.

ITC are reactive electrophiles that have been studied in the context of a broad range of pharmacological effects using cell-based assays and *in vivo* model systems. For instance, the ITC sulforaphane, derived from hydrolysis of glucoraphanin, is an established inducer of nuclear factor (erythroid-derived 2)-like 2 (Nrf2). Nrf2 activation by sulforaphane upregulates xenobiotic metabolizing and antioxidant-responsive phase II enzymes like NAD(P)H quinone oxidoreductases and glutathione S-transferases (Traka and Mithen, 2009). Beyond xenobiotic metabolism, Nrf2 is a repressor of genes involved in hepatic lipid synthesis (Vomhof-DeKrey and Picklo, 2012), consistent with data from human intervention studies showing that Beneforte broccoli also reduces plasma LDL-cholesterol to a greater extent than conventional broccoli (Armah et al., 2015). In addition to modulating plasma metabolite profiles and contributing to chemoprotection, sulforaphane has been studied in the context of diabetes, with sulforaphane treatment resulting in decreased glucose production in rat hepatoma cell lines, mediated in part by Nrf2 downregulation of gluconeogenesis, and reduced fasting blood glucose levels in human patients with Type 2 diabetes (Axelsson et al., 2017). Beyond the physiological impacts associated with the induction of Nrf2, ITC have also been shown to interact with sensory and inflammatory pathways. Allyl ITC, derived from the GS sinigrin, is an agonist of the transient receptor potential ion channel TRPA1, which is involved in signaling inflammatory pain (Bellono et al., 2017), while sulforaphane has been shown to covalently modify lipopolysaccharide-activated Toll-like receptor 4 (TLR4), resulting in the reduced secretion of pro-inflammatory cytokines in human peripheral blood mononuclear cells and monocytes (Folkard et al., 2014).

Plant and gut bacterial metabolism have been identified as two different routes for the conversion of GS into ITC products required for biological activity. Like many phytonutrients relevant to human health, GS in crucifers are involved in chemical defense *in planta* (Halkier and Gershenzon, 2006). These metabolites are stored in the plant tissue in an inert, pro-drug-like state and are converted into an active state by myrosinase enzymes, thio-specific glucoside hydrolases found in cruciferous vegetables (Figure 1A). GS are sequestered in vacuoles under normal conditions, while myrosinases are expressed in distinct, specialized myrosin cells (Kissen et al., 2009). Upon disruption of the plant tissue, myrosinases mediate the hydrolysis of GS into a variety of possible products, including ITC, nitriles, and epithionitriles, determined by environmental pH and the presence of specifier proteins.

In addition to being activated by native plant enzymes, GS can also be hydrolyzed to ITC through the action of the intestinal microbiota (Rabot et al., 1993). This route of GS activation plays a particularly important role in a dietary context, since plant myrosinases are often denatured during cooking. Consumption of cooked broccoli by human volunteers following a regimen to reduce intestinal colonization through antibiotic treatment and mechanical cleansing resulted in significantly diminished quantities of excreted ITC-derived metabolites compared to volunteers who did not follow the microflora-reducing regimen (Shapiro et al., 1998). An interesting consequence of the role that the gut microbiota plays in activating these prodrug-like molecules is significant inter-individual variability in the level of bioactive ITCs produced following a meal containing cooked cruciferous vegetables (Navarro et al., 2014; Shapiro et al., 2001). The observed variability has been shown to be associated in part with the diversity of metabolic capabilities represented in the intestinal microbiota, with *ex vivo* fecal bacterial cultures from high ITC-excreting individuals turning over larger quantities of glucosinolate than those from low ITC-excreting individuals (Li et al., 2011). Several gut bacterial are known to metabolize GS, including strains of *Lactobacillus*, *Bifidobacterium*, and *Bacteroides* (Cheng et al., 2004; Elfoul et al., 2001; Palop et al., 1995). As is the case with many of the interactions between dietary plant metabolites and the gut microflora, however, the bacterial genes encoding this conversion have not been identified.

Here, we investigate the genetic and biochemical basis behind the activation of dietary GS by *Bacteroides thetaiotaomicron* (*Bt*), a prominent member of the human gut microbiota that has been the subject of extensive research (Xu et al., 2003). Using insertional mutagenesis, we identified a previously uncharacterized carbohydrate-responsive operon, *BT2160-BT2156*, required for GS conversion to ITC in *Bt*. While microbial genetic analysis showed that *BT2157* and *BT2158* are needed for activity in *Bt*, biochemical analyses revealed that the coordinated action of BT2158 with either BT2156 or BT2157 can promote GS transformation *in vitro*. Furthermore, mice monocolonized with mutant *Bt* lacking the complete operon (*Bt*Δ*2157*) showed decreased levels of isothiocyanate production when fed glucosinolates. Collectively, these results increase our resolution of gut microbial processing of plant metabolites, a first step towards quantifying and predicting the pharmacological impacts that these dietary plant molecules have on disease prevention and progression.

## RESULTS

### Glucosinolate metabolizing activity varies among *Bacteroides thetaiotaomicron* strains

We began our investigations by validating the ability of a panel of *Bt* strains to convert GS to ITC. We cultured human-associated isolates *Bt* VPI-5482, *Bt* 8736, *Bt* 7330, and *Bt* 3731 under standard anaerobic growth conditions in rich medium containing glucotropaeolin (BGS), a commercially available GS with a benzyl moiety found in garden cress. Culturing medium was supplemented with cysteine, included as a reducing agent (Bacic and Smith, 2008) and for *in situ*, reversible capture of any formed ITC (Angelino et al., 2015; Baillie and Slatter, 1991; Bruggeman et al., 1986). Conjugates of benzyl isothiocyanate with cysteine (BITC-cys) are readily detectable using liquid chromatography-mass spectrometry (LC-MS) and were measured in the spent media (Figure 1B). The surveyed *Bt* strains displayed variable metabolic activity in rich media containing glucose (Figure 1C), revealing that different levels of glucosinolate- and non-glucosinolate-metabolizing *Bt* strains may be a potential source for inter-individual variation in ITC production in human feeding studies. Other products of BGS metabolism, such as nitriles or epithionitriles, were not detected by LC-MS. To confirm that the hydrolysis of GS to ITC does not occur spontaneously in rich medium, including under the low pH conditions induced by *Bt* growth, we incubated BGS in sterile-filtered spent media from a wild-type *Bt* VPI-5482 culture and observed that no BITC-cys was formed (Figure S1A). This result also suggests that the proteins responsible for GS metabolism are cell-associated.

### A loss of function library screen yields an operon necessary for glucosinolate conversion to isothiocyanate in Bt

Because BLAST analysis revealed no *Bt* homologs to characterized plant myrosinases, we sought to identify candidate GS metabolizing genes in *Bt* with an untargeted genome-wide screen. We used the Mariner transposon to generate an insertion library in *Bt* VPI-5482 (Goodman et al., 2009) and manually selected ~7500 clones for screening using a coupled growth assay. While growth of *Bt* is not inhibited by ITC production (Figure S2A), ITC exhibits bactericidal and bacteriostatic activity towards some bacterial strains (Dufour et al., 2015). We used this observation to develop a coupled growth assay that links ITC production by *Bt* transposon mutants to growth inhibition of an ITC-sensitive *Escherichia coli* (*E. coli*) strain inoculated into spent media from *Bt* cultures grown with GS (Figure S2B). *Bt* mutants yielding spent media that did not inhibit *E. coli* growth were selected for further evaluation of transposon insertions disrupting candidate ITC-producing genes (Figure S2C). In order to avoid false positives from transposon mutants that hinder or abolish *Bt* growth, leading to low ITC production and low resulting toxicity to *E. coli*, we monitored *Bt* growth by optical density (OD) measurements and disregarded *Bt* mutants with low growth (Figure S2D).

We next used semi-random PCR to identify the genes hit in the screen (Goodman et al., 2009), resulting in a list of 26 genes with several loci hit multiple times (Table S1). While several candidates emerged from this screen, the most highly represented hits contained insertions in two gene clusters, predicted to be operons. The most abundantly represented operon, *BT1220-BT1222*, contains genes involved in the pentose phosphate pathway. Because these genes encode a pathway in central carbon metabolism, we considered them unlikely to be directly responsible for the enzymatic conversion of GS to ITC and excluded them from further evaluation as potential GS metabolizing enzymes.

The second most abundantly represented operon, *BT2159-BT2156*, included genes with predicted carbohydrate metabolizing activity and became our primary candidate for thioglucosidase function. The putative annotations ascribed to the genes in this operon include two nicotinamide-dependent oxidoreductases (BT2158 and BT2159), a glycoside hydrolase (BT2157), and a sugar-phosphate epimerase/isomerase (BT2156), under the control of transcriptional regulator BT2160 (Figure 2A). *In silico* analysis of the encoded protein sequences predicted that BT2160 spans the inner membrane, BT2159 is cytoplasmic, BT2158 is periplasmic, and BT2157 is an outer membrane lipoprotein (Figure S3A) (Käll et al., 2007; Petersen et al., 2011; Yu et al., 2010). While BT2156 contains a signal peptide sequence and is predicted to be non-cytoplasmic, the localization could not be more specifically predicted.

**Figure 2.**
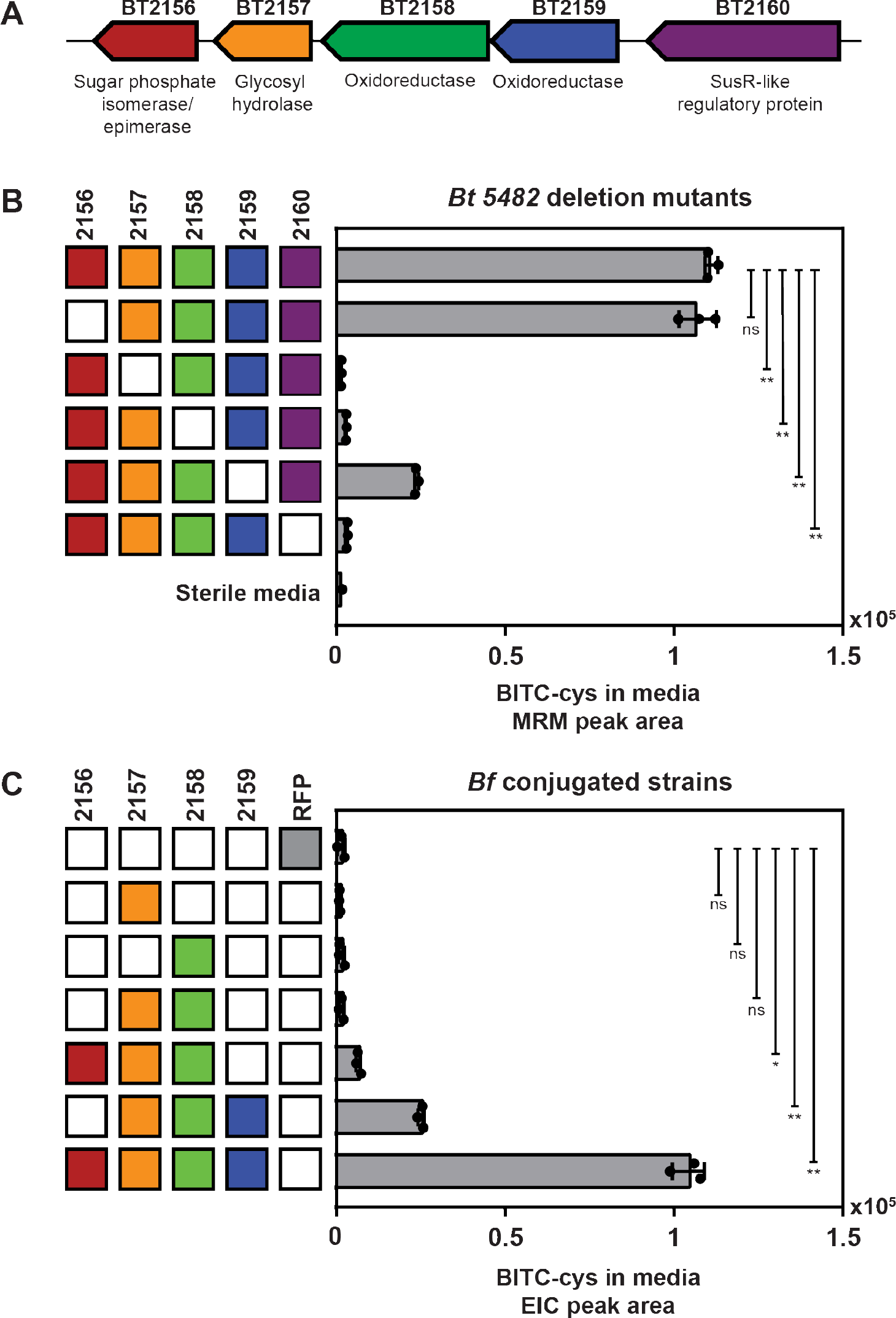
An operon necessary for glucosinolate conversion to isothiocyanate in *Bt*. (A) The operon identified to be involved in glucosinolate metabolism in *B. thetaiotaomicron* VPI-5482. Predicted functions based on homology are annotated below each gene. See also Figures S1, S2, and S3, and Table S1. (B) BITC-cys produced from BGS by *Bt* mutants with single deletions in the BT2160-2156 operon grown in rich glucose media for 48 hours. Filled boxes represent natively expressed genes, while empty boxes represent deleted genes. Multiple reaction monitoring (MRM) by LC-MS/MS was used to track the transition of the protonated BITC-cys molecule with *m/z* 271.0 to a product ion with *m/z* 122.0. Data shown is the mean±SD of three biological replicates, with individual replicates shown as filled circles. BITC-cys background in sterile media is shown with one replicate on the bottommost row. Significant differences from the wild-type *Bt* mean are marked by ns (not significant) or **(p<0.0001), as determined using Dunnett’s multiple comparison test. See also Figure S1. (C) BITC-cys produced from BGS by strains of *Bf* expressing subsets of the BT2160-2156 operon after 48 hours of culturing in rich glucose media. Filled boxes represent extra-chromosomally complemented genes. Extracted ion count (*m/z* 271.0570) with LC-MS was used to track quantities of protonated BITC-cys molecule. Data shown is the mean±SD of three biological replicates, with individual replicates shown as filled circles. Significant differences from the RFP expressing negative control strain are marked by ns (not significant), * (p<0.05), or **(p<0.0001) as determined using Dunnett’s multiple comparison test. See also Figure S1.

This operon is one of many abundant carbohydrate metabolizing loci in *Bt*, including 261 glycoside hydrolases and polysaccharide lyases, thought to be involved in the ability of *Bt* to utilize a diverse spectrum of plant glycans (Martens et al., 2009). To assess how widely the *BT2159-BT2156* operon is represented across various bacterial strains, we performed a BLAST search for homologs with greater than 60% identity on the amino acid level and greater than 60% coverage of the query sequence. The results of this analysis revealed that this operon is highly conserved across various *Bacteroides* species, including strains of *Bacteroides caccae* and *Bacteroides ovatus* (Figure S3B). Recent chemical genomics studies have identified this operon as being important for catabolism of the disaccharides trehalose, leucrose, palatinose, and raffinose (Liu et al., 2019). Gene expression data from chemostat-grown monocultures of *Bt* (GEO dataset GPL1821) showed that the operon genes are strongly upregulated by glucose-containing media relative to pig mucin glycan (PMG)-containing media (Sonnenburg et al., 2006). Disruption of this operon also lowered fitness of *Bt* in mice when *Bt* was co-colonized with *Eubacterium rectale* or with a community of Firmicutes but not when *Bt* was used in mono-colonization or in a community of other *Bacteroides* strains (Goodman et al., 2009). Despite their abundant representation in *Bacteroides* genomes and their importance for *Bt* fitness, however, none of these proteins and or their close homologs have been biochemically characterized.

### Two enzymes are necessary for GS transformation in *Bt*

To identify which genes in the *BT2159-BT2156* operon are directly involved in GS hydrolysis, we generated targeted genomic deletions. Each open reading frame in the operon was replaced with an in-frame start-stop codon pair using the pExchange recombination system (Koropatkin et al., 2008). We found the growth of the knockout strains to be consistent with wild-type *Bt* growth in rich media containing glucose (Figure S1C). Deletion mutants were cultured in rich medium supplemented with BGS and assayed for BITC-cys in spent media using LC-MS/MS (Figure 2B). As anticipated, knockout of the putative transcriptional regulator BT2160 resulted in loss in GS metabolizing activity. Of the genes that encode enzymes, *BT2157* was necessary for ITC production in *B*t and deletion of *BT2158* resulted in only trace levels of BITC-cys production. Deletion of *BT2159* was also observed to be detrimental to GS metabolism but detectable levels of ITC were still produced, while knockout of BT2156 did not significantly affect glucosinolate metabolism in *Bt*.

From simultaneous time course measurements of wild-type *Bt* growth and BITC production in rich medium containing glucose, we observed that ITC accumulation and GS depletion became significant only in late stages of growth (Figure S1B). This result suggested that GS was being used as a late-stage carbon source following depletion of glucose in the media. To ensure that the *Bt* mutants maintained consistent GS metabolizing behavior across multiple carbon sources, we cultured the *Bt*Δ*2157* strain in rich medium supplemented with different carbohydrates and BGS. In each of the six carbon sources surveyed, *Bt*Δ*2157* lacked the ability convert GS to ITC (Figure S1E).

To support the role of BT2157 and BT2158 in the conversion of GS to ITC in *Bt*, we used the pFD340 plasmid-based expression system (Smith et al., 1992) to complement single deletion mutants *Bt*Δ*2157* and *Bt*Δ*2158* with their respective deleted genes. Constitutive expression of the corresponding deleted gene rescued GS metabolizing activity for both strains, confirming that ITC production is only observed when both BT2157 and BT2158 are present in *Bt* (Figure S1F).

### Expression of the BT2159-BT2156 operon in an inactive *Bacteroides* species results in gain of activity

We sought to determine if the BT2157 and BT2158 enzyme pair represents the minimal unit required for GS hydrolysis in the heterologous host *Bacteroides fragilis* (*Bf*), which we determined is unable to metabolize GS present in culture media (Figure 1C). BLAST analysis also established that the *Bf* genome contains no close homologs of *BT2160-BT2156*. Using the pFD340 expression system (Smith et al., 1992), we expressed subsets of enzymes encoded by the *BT2159-BT2156* operon in *Bf*, cultured these strains in rich medium supplemented with BGS, and assayed for BITC-cys in spent media using LC-MS/MS. Growth of these *Bf* strains was also monitored to ensure that the constitutive expression of these genes did not result in growth defects relative to a control strain expressing RFP (Figure S1D). We found that simultaneous expression of BT2157 and BT2158 was not sufficient for gain of GS metabolizing activity relative to the RFP negative control (Figure 2C). In contrast, expression of either BT2156 or BT2159 in addition to BT2157 and BT2158 resulted in measurable quantities of ITC production. Furthermore, expression of all four enzymes in the operon results in gain of GS metabolizing function in *Bf*.

### *In vitro* conversion of glucosinolate to isothiocyanate requires BT2158 and either BT2156 or BT2157

With the observation that BT2157 and BT2158 are important but not sufficient for activity *in vivo*, we next aimed to test the function of these proteins *in vitro* to better understand which of these enzymes act directly on GS. We used *E. coli* to individually express and purify 6x His-tagged variants of each of the proteins in the *BT2159-BT2156* operon. While three of the proteins were expressed with sufficient yields to permit *in vitro* assays, we achieved only low yields of BT2158 expression (Figure S4A). In an effort to improve the yield of soluble protein, we turned to *Bt* as an expression host. We used a *Bf* phage promoter sequence to drive overexpression of His-tagged BT2158 with its native ribosome binding site in the *Bt*Δ2158 mutant (Whitaker et al., 2017). This approach resulted in increased levels of soluble target protein (Figure S4B). To ensure that native *Bt* proteins were not co-eluting or being pulled down with the His-tagged BT2158, we analyzed the purified protein fraction with SDS-PAGE and silver staining and observed only a single protein band corresponding to the mass of BT2158 (Figure S4C).

Before initiating biochemical assays, we also considered the cofactors that might be required for protein activity. InterPro analysis revealed the presence of domains in BT2159 and BT2158 unique to the MocA/Idh/Gfo family of proteins (Figure S3A), members of which are known to bind NAD(P) (Taberman et al., 2016). Members of this protein family include the myo-inositol 2-dehydrogenase from *Bacillus subtilis* (BsIDH) and the aldose oxidoreductase from *Caulobacter crescentus*. Protein sequence alignment of BT2158 with BsIDH showed that residues Lys-133, Asp-248, and His-252 correspond to Lys-97, Asp-172, and His-176 of BsIDH, conserved residues that form a triad involved in hydride transfer (van Straaten et al., 2010). Based on this analysis, we chose to include NAD^+^ in our initial experiments.

Having expressed and purified the complete set of operon proteins, we next incubated different combinations of purified BT2159-BT2156 in the presence of BGS and NAD in order to identify the minimal set of proteins required for *in vitro* GS conversion. After quenching the reactions with acetonitrile and addition of cysteine for ITC conjugation, we measured BITC-cys formation and BGS depletion using LC-MS (Figure 3A). As with the complemented *Bf* strains, individual proteins alone did not mediate BGS conversion *in vitro*. When pairs of proteins were tested, however, measurable activity was observed for BT2158 when combined with either BT2156 or BT2157. In contrast to the *in vivo* analysis of *Bf* strains expressing operon proteins, however, addition of BT2156 and BT2159 to BT2157 and BT2158, representing the complete set of proteins encoded by the operon, did not seem to enhance BGS turnover. Taken together, our data indicates that BT2158 is necessary but not sufficient for glucotropaeolin hydrolysis *in vitro*.

**Figure 3.**
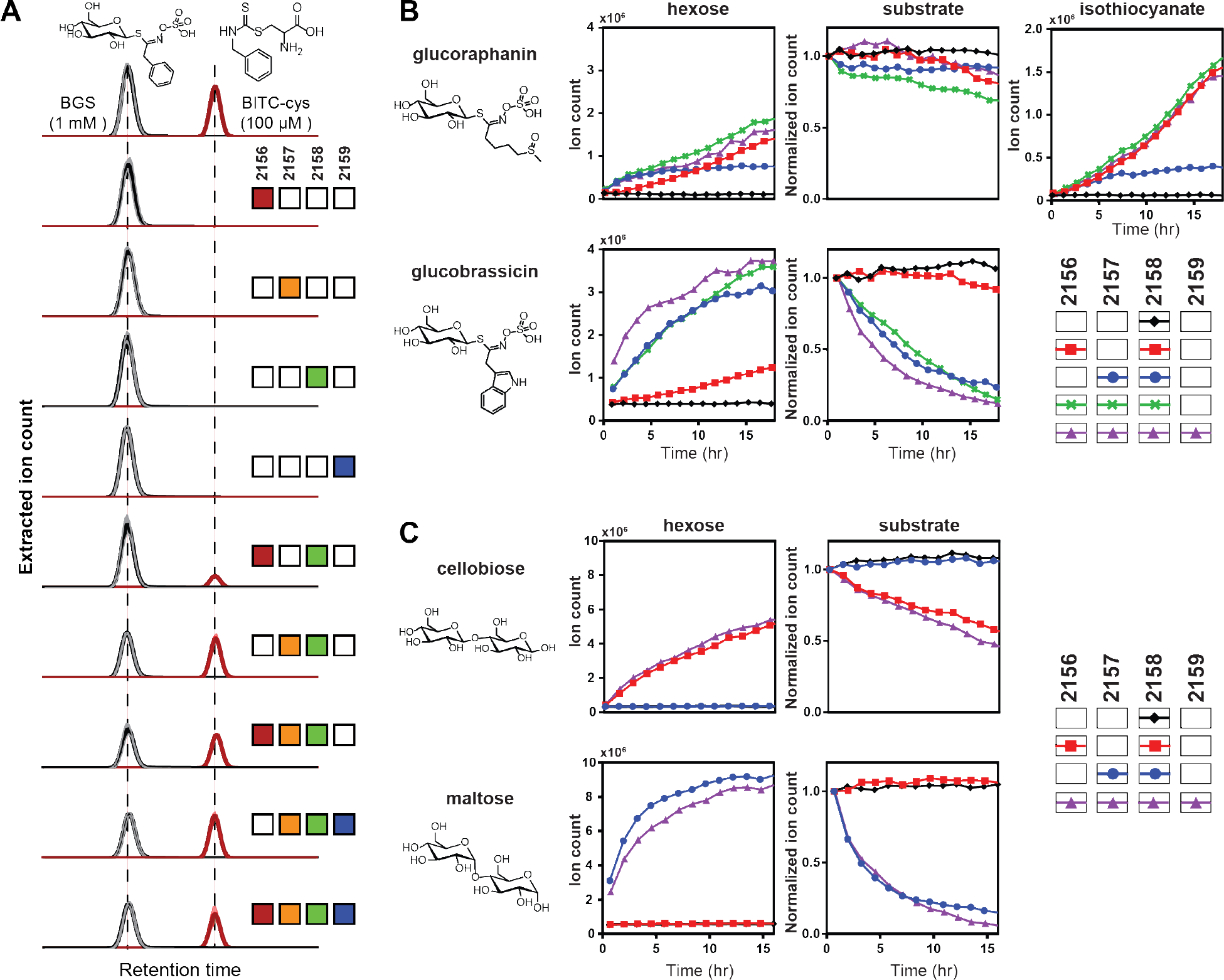
*In vitro* glucosinolate conversion by recombinant proteins. (A) Extracted ion chromatograms of glucotropaeolin (black; *m/z* 408.0428) and BITC-cys (red; *m/z* 271.0570) produced by different combinations of BT2159-BT2156 proteins *in vitro*, represented on the same scale. BT2158 was expressed in and purified from *Bt*Δ2158 and all other proteins were expressed in and purified from *E.coli* BL21 DE3. The first row of chromatograms represents 1 mM glucotropaeolin and 100 μM BITC-cys standards. Data shown represent the mean of three replicates, with the shaded regions representing one standard deviation. See also Figure S4. (B) Direct injection MS extracted ion counts showing glucose release (*m/z* 179.0561), substrate consumption (glucoraphanin: *m/z* 436.0411; glucobrassicin: *m/z* 447.0538), and isothiocyanate (sulforaphane: *m/z* 178.0355) production over time by different combinations of BT2159-BT2156 proteins acting on glucoraphanin and glucobrassicin. No corresponding isothiocyanate was observed for glucobrassicin. Substrate ion count was normalized against the initial ion count for each combination of proteins. Data shown is one replicate, representative of triplicate results. (C) Direct injection MS extracted ion counts showing glucose release (*m/z* 179.0561) and substrate consumption (*m/z* 341.1089) over time by different combinations of BT2159-BT2156 proteins with cellobiose and maltose. Substrate ion count was normalized against the initial ion count for each combination of proteins. Data shown is one replicate, representative of triplicate results.

In order to further investigate the cofactor requirements of the *Bt* proteins, we tested BT2157 and BT2158 with a panel of nicotinamide cofactors, using LC-MS to quantify BITC-cys formation (Figure S4D). While the presence of any of the four nicotinamide cofactors improved BGS conversion by BT2157 and BT2158, NAD^+^ resulted in the most significant increase in activity by the protein pair. However, basal levels of activity were still observed in the absence of added cofactor, suggesting either that the nicotinamide cofactor is not necessary for activity or that low levels of cofactor were co-purified with the proteins.

We also tested the activity of BT2159-BT2156 towards glucosinolates with side groups containing non-aromatic moieties, including glucoraphanin (GRP), which has been directly associated with the chemopreventive effects of brassicas (Jeffery and Araya, 2009), and glucobrassicin (GBR), another dietarily relevant glucosinolate found in broccoli (Figure 3B). GS conversion and glucose production over the course of the reaction were tracked by hourly direct injections with time-of-flight mass spectrometry. While sulforaphane production from glucoraphanin was also simultaneously measured, mass features corresponding to the ITC derived from GBR were not observed, likely due to the instability of indole isothiocyanates (Agerbirk et al., 2009). As observed with BGS, BT2158 with either BT2156 or BT2157 represent the minimal set of proteins required for activity on GRP or GBR. In conjunction with the evidence from the knockout *Bt* strains and complemented *Bf* strains, these *in vitro* results indicate that BT2158 is necessary for GS hydrolysis to ITC, but that the addition of other operon proteins, namely BT2156 or BT2157, is required for activity.

### BT2159-BT2156 metabolize a selection of disaccharides *in vitro*

As suggested by the predicted carbohydrate-metabolizing annotations ascribed to the *BT2160-BT2156* operon, we considered that other dietarily abundant carbohydrates might also serve as substrates. To explore this hypothesis, we incubated different combinations of purified protein with NAD^+^ and a panel of dietary disaccharides, and measured substrate consumption and hexose production over the course of the reaction by direct injection mass spectrometry. Subsets of the operon proteins were found to hydrolyze both cellobiose and maltose, with BT2156 and BT2158 representing the minimal set required for activity on cellobiose, and the combination of BT2157 and BT2158 required for activity on maltose (Figure 3C). In contrast to the redundant activity we observed with different pairs of enzymes on GS substrates, it is notable that each of these carbohydrate substrates was acted on by only one pair of proteins. The pair of BT2157 and BT2158 did not exhibit activity on cellobiose and BT2156 with BT2158 did not act on maltose. Maltose and cellobiose are both composed of two glucose units linked by an alpha- or beta-glycosidic linkage, respectively, suggesting that BT2157 or BT2156 may dictate α-versus β-O-glucosidase activity when combined with BT2158. For both disaccharides, addition of the remaining operon proteins did not significantly improve activity. Carbohydrates that were not significantly hydrolysable by any protein combination tested include raffinose and lactose (Figure S4E), potentially indicating that a substrate with glucose on the non-reducible end is required for activity.

### Isothiocyanate production is reduced in gnotobiotic mice colonized with a *Bt* mutant deficient in glucosinolate metabolism

To determine if this locus is required for GS processing by *Bt* in the context of gut colonization, we tested GS metabolism in germ-free mice mono-colonized with wild-type *Bt* or *Bt*Δ2157 (Figure 4A). Mice were fed *ad libitum* a polysaccharide-deficient chow diet, supplemented with two consecutive daily doses of pure BGS by gavage. We observed higher levels of BITC N-acetyl cysteine and BITC-cys, established biomarkers of crucifer intake and ITC exposure (Hwang and Jeffery, 2003), in the urine of wild-type *Bt-*colonized mice compared to *Bt*Δ2157-colonized or germ-free mice (Figure S5C). These data provided initial evidence that the BT2159-BT2156 operon plays a role in the metabolism of GS to ITC *in vivo*, corroborating our genetic and *in vitro* biochemical findings. Although we observed higher levels of BITC-derived metabolites in the urine and feces of *Bt-*colonized mice following BGS gavage, this effect was inconsistent; moreover, we detected some background GS hydrolysis in germ-free mice, as was noted in a previous report (Budnowski et al., 2015).

**Figure 4.**
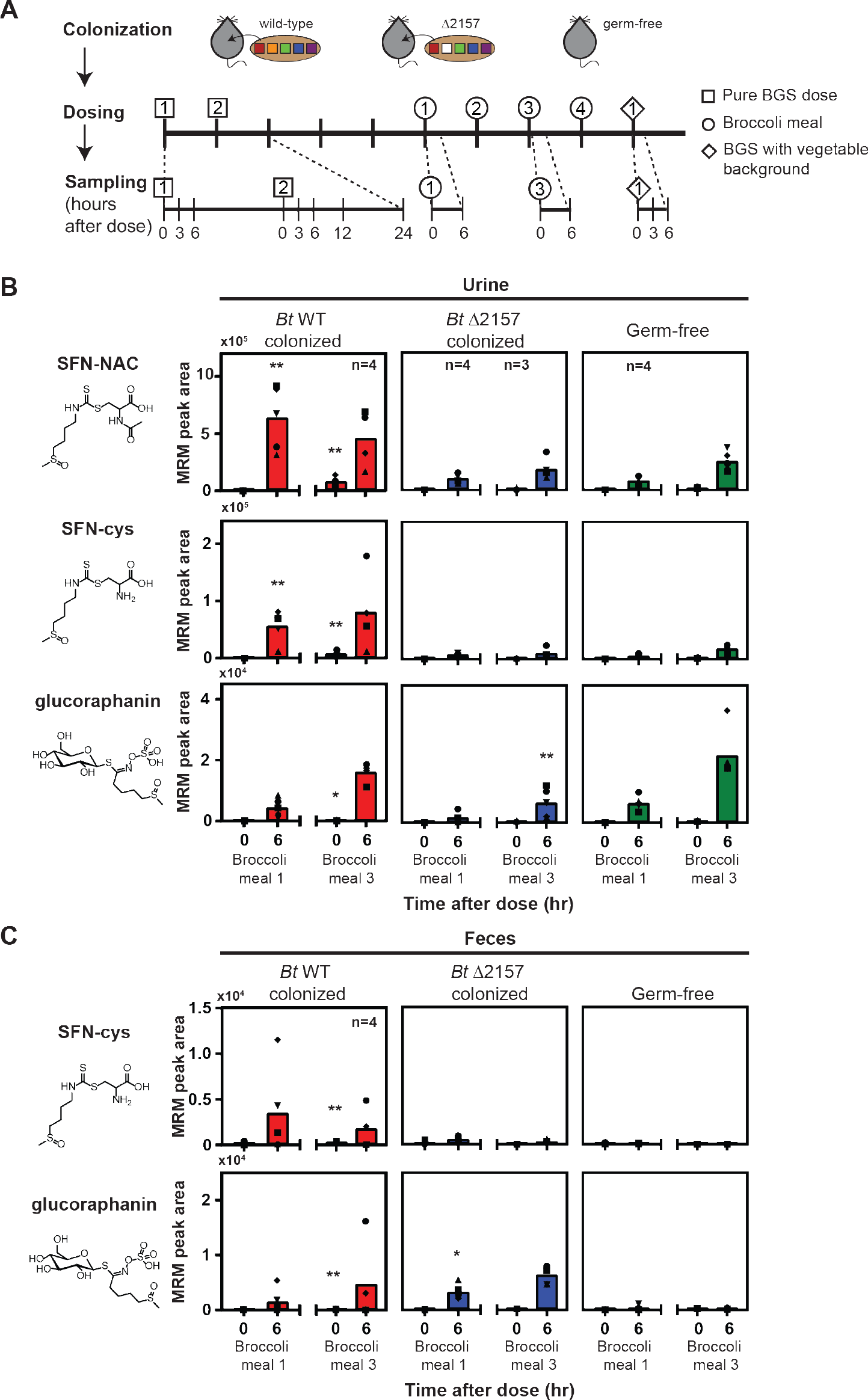
Monocolonization of gnotobiotic mice by mutant *Bt*. (A) Schematic for colonization, dosing schedule, and collection of urine and feces. Each mark in the dosing schedule corresponds to 24 hours. Numbers in the unfilled shapes represent the dose number within that series. (B) SFN-NAC and SFN-cys mercapturic acid conjugates and unconverted glucoraphanin excreted in urine six hours after doses of broccoli meal containing glucoraphanin. See also Figure S5. (C) SFN-cys and unconverted glucoraphanin in feces collected six hours after doses of broccoli meal containing glucoraphanin. SFN-cys was the product of *in situ* conjugation of free SFN with exogenously applied cysteine. Bars represent the mean values for each group, with individual data overlaid (different shapes correspond to individual mice). Metabolites were measured by LC-MS/MS and quantified by MRM (SFN-NAC: *m/z* 341.1→ *m/z* 177.9; SFN-cys: *m/z* 299.1→ *m/z* 136; glucoraphanin: *m/z* 436.0→ *m/z* 372). n=5 for each colonized group and time point unless otherwise noted on the graph. ** represents significance with p<0.01 from the germ-free control at the corresponding time point. * represents significance with p<0.05. Statistical groups were determined using the Tukey test.

Reasoning that both issues might stem from administration of pure glucosinolate outside of the context of a food matrix, we administered glucosinolate in the form of reconstituted broccoli meals by gavage, followed by one bolus of a GS-free broccoli meal supplemented with pure BGS the following week (Figure 4A). Though broccoli meals were found to contain a variety of glucosinolates including neoglucobrassicin, hydroxyglucobrassicin, and erucin (Figure S5B), we limited our analysis of urine and fecal samples to glucotropaeolin and glucoraphanin, along with their respective mercapturic acid ITC conjugates, due to the established biological activities of their corresponding isothiocyanates. Six hours after the broccoli meals, higher levels of SFN-NAC and SFN-cys were consistently observed in the urine and feces, respectively, of mice colonized by *Bt* versus *Bt*Δ*2157* (Figure 4B). Consistent with our observations with pure compound feeding, higher levels of SFN-cys, the upstream metabolite of SFN-NAC in the mercapturic acid pathway, were also observed in the urine of mice colonized with wild-type *Bt* compared to *Bt*Δ*2157* or germ-free mice. Taken together, these experiments indicate that the *BT2159-BT2156* operon plays an important role in glucosinolate hydrolysis *in vivo*.

## DISCUSSION

Despite the established impact of dietary plant metabolites on human health and the role of the gut microbiome in modulating their activity, few genes from the gut microbiota required for conversion and, in many cases, activation of these molecules have been described. Our understanding of the observed metabolism has often been limited to identification of the involved bacterial strains (Clavel et al., 2006; Tsuchihashi et al., 2008); for example, the soy isoflavone daidzein (Bowey et al., 2003) and the stilbenoid resveratrol found in grapes (Bode et al., 2013) both undergo microbiome-mediated processing. One of the few studies that report specific bacterial genes and mechanisms responsible for metabolism of bioactive plant molecules is the identification of a cytochrome-encoding operon responsible for inactivation of the plant-derived cardiac drug digoxin by the gut bacterium *Eggerthella lenta* (Haiser et al., 2013). The lack of knowledge regarding the genetic basis for production of reactive isothiocyanates by gut microbiota limits our ability to understand inter-individual differences in GS hydrolysis and use quantitative approaches to connect isothiocyanate levels to host effects.

Our efforts to identify a genetic basis for glucosinolate conversion to isothiocyanates by the human gut microbiota have led to the identification of an operon in *Bt* that is necessary and sufficient for glucosinolate activation and is also able to metabolize selected disaccharides. To date, identified bacterial glycosidases able to cleave the thio-linked sugar of glucosinolates include a glycoside hydrolase family 3 β-O-glucosidase in the soil-isolate *Citrobacter* WyE1(Albaser et al., 2016); after determining that cell-free protein extract was capable of degrading glucosinolates, activity-guided purification was performed to isolate the active protein. A 6-phospho-β-glucosidase was also determined to be associated with glucosinolate metabolism in the pathogenic *E. coli* strain 0157:H7, identified through homology to characterized plant myrosinases (Cordeiro et al., 2015); however, deletion of the homologous genes did not abolish the ability of the bacteria to hydrolyze the aliphatic GS sinigrin. While, to our knowledge, there have not been any reports to date of bacterial myrosinase-encoding genes that are both necessary and sufficient for GS hydrolysis, the *sax* operon in certain pathovars of the plant pathogen *Pseudomonas syringae* has been identified as being necessary and sufficient for overcoming aliphatic isothiocyanate toxicity (Fan et al., 2011). Putative myrosinase activity by the 6-phospho-β-glucosidase from *E. coli* 0157:H7, in conjunction with sinigrin-induced activation of a glucose phosphotransferase system in *E. coli* VL8, has led to the proposal that phosphorylation of the glucose moiety may be prerequisite for glucosinolate hydrolysis in these bacterial strains (Narbad and Rossiter, 2018). While the annotation of BT2156 as a putative sugar phosphate epimerase/isomerase may suggest that BT2159-BT2156 act on a similarly phosphorylated substrate, our kinetic *in vitro* reactions using purified proteins were performed in phosphate-free buffers and in the absence of externally supplied ATP (Figures 3B **and** C). In addition, features with masses corresponding to phosphorylated glucosinolates or phosphorylated hexose were not detected in either *in vitro* reactions or the spent media of *Bt* strains.

While Bacteroidetes are well known metabolizers of a broad spectrum of plant glycans, the question remains whether the *BT2160-BT2156* operon was evolved to specifically process glucosinolates. *Bacteroides* spp. have been found to activate a variety of plant metabolites, including the lignin secoisolariciresinol diglucoside (SDG) (Clavel et al., 2006) and the flavonoid glycoside rutin (Bokkenheuser et al., 1987). The β-glucosidase involved in SDG hydrolysis in *Bacteroides uniformis* ZL1 was also found to exhibit activity on other plant glycosides, including astragalin, rutin, and isoquercetin (Tao et al., 2014). Similarly, an α-L-rhamnosidase from *Bacteroides* JY-6 was isolated for its activity on different rhamnoglucosides, including rutin, poncirin, and hesperidin (Jang and Kim, 1996). BT2159-BT2156 exhibits similarly broad activity towards the disaccharides cellobiose and maltose *in vitro*, which suggests that this operon may have evolved to process other glycosides in addition to glucosinolates. The native substrate or function of the proteins encoded by this operon, however, is yet to be determined.

Of the 88 PULs in *Bt*, the starch-utilization system (Sus) is the most well-characterized and is comprised of eight proteins that work in a cell-associated, concerted manner to bind and degrade starch (Martens et al., 2009). The Sus system is regulated by the membrane protein SusR, which is activated upon binding with maltose or higher oligomers. Efforts to identify similarly regulated loci have revealed four SusR paralogs in *Bt*, including BT2160 (Ravcheev et al., 2013). Though BT2160 is annotated as a SusR-like protein, there are a few differences between Sus and BT2160-BT2156. BT2160-BT2156 is not classified as a PUL because it lacks homologs to SusC or SusD, involved in starch binding and transport (Foley et al., 2016). Our *in vitro* studies with purified protein suggest that coordinated action by the periplasmic BT2158 with either BT2156 or BT2157 is required for hydrolytic activity. In contrast, SusA and SusB are involved in hydrolytic activity in the periplasm, but do not require coordinated activity with another enzyme. In addition, individual knockouts of SusA or SusB were not deleterious to growth on starch (D’Elia and Salyers, 1996), whereas single knockouts of BT2157 and BT2158 both resulted in a loss of GS conversion. SusG is annotated as an outer membrane-bound glycosyl hydrolase and was found to be necessary but by itself insufficient for growth on starch, similarly to BT2157 (Shipman et al., 1999). In the starch utilization system, outer membrane proteins SusC-G form a complex that is required for starch binding and transport. SusC and SusD individually are insufficient for starch binding, but together are capable of binding 60% of the starch bound by wild-type *Bt*, with the addition of SusE and SusF providing the remaining starch affinity (Shipman et al., 2000). Though BT2157 and BT2158 together are sufficient for GS hydrolysis *in vitro*, expression of this protein pair in the inactive *Bf* did not result in gain of activity. Additional expression of BT2156 or BT2159 increased activity to 6% and 24%, respectively, of the activity of the *Bf* strain expressing the full operon, suggestive of the possibility that these proteins may similarly require formation of a complex for GS hydrolyzing activity. Interestingly, the *in vitro* hydrolysis of cellobiose and maltose require different combinations of proteins. While the mechanism behind glucosinolate activation and disaccharide cleavage by these proteins is yet to be determined, the different combinations of proteins required for different substrates may provide insight into the catalytic roles that each of the proteins required for activity plays in hydrolysis.

In this study, we have identified a genetic and biochemical basis for the activation of a class of dietary plant metabolite, the glucosinolates, by a widely conserved bacterial species in the human gut microbiota. Glucosinolates have been extensively studied for the role of the activated product, isothiocyanates, on human health. Despite their importance, genes in the gut microbiota that contribute to activation have remained elusive. Using a genome-wide screen, we identified the operon *BT2159-BT2156* as a key player in glucosinolate conversion to isothiocyanates by *Bt*. Discovery of the microbial pathways that produce bioactive compounds such as isothiocyanates, in conjunction with recent developments in genetic engineering tools for cellular therapeutics (Mimee et al., 2015; Whitaker et al., 2017) can contribute toward engineering gut commensals for therapeutic benefit. On a more fundamental level, however, this work provides a critical advance in our understanding of the many interrelated mechanisms through which the gut microbiota and plant-derived metabolites in our diet interact to influence health and disease.

## ACKNOWLEDGEMENTS

This work was supported by an HHMI and Simons Foundation Grant 55108565 and NIH DP2AT00832101 (to E.S.S.), a Stanford Bio-X Bowes (to C.S.L.), and ChEM-H seed grant funding (to J.L.S. and E.S.S.). We thank M. Fischbach, E. Carlson, and M. Voges (Stanford) for comments on the manuscript, and K. Smith and other members of the Sattely lab for helpful discussions. We thank K. Ng, D. Dodd, and E. Shepherd (Stanford) for assistance with and advice on gnotobiotic mouse studies, and W. Whitaker (Stanford) for providing the *Bacteroides* phage promoter sequence. We thank C. Miller, S. Hull, and S. Lensch (Stanford) for their efforts on *in vitro* biochemical characterization. We thank S. Mazmanian (California Institute of Technology) for providing plasmid pFD340. We thank the Stanford High-Throughput Bioscience Center for technical assistance with the *Bt* mutant screen.

## CONTRIBUTIONS

C.S.L., S.J.S., C.A.C.D., A.P.K., C.R.F, J.L.S., and E.S.S designed experiments and analyzed data. C.S.L., S.J.S., C.A.C.D., A.P.K, C.R.F., and S.K.H. performed experiments. C.S.L. and S.J.S. wrote the paper with edits from E.S.S., C.A.C.D., A.P.K., and C.R.F.

## DECLARATION OF INTERESTS

The authors declare no competing interests.

## SUPPLEMENTAL INFORMATION

**Figure S1.**
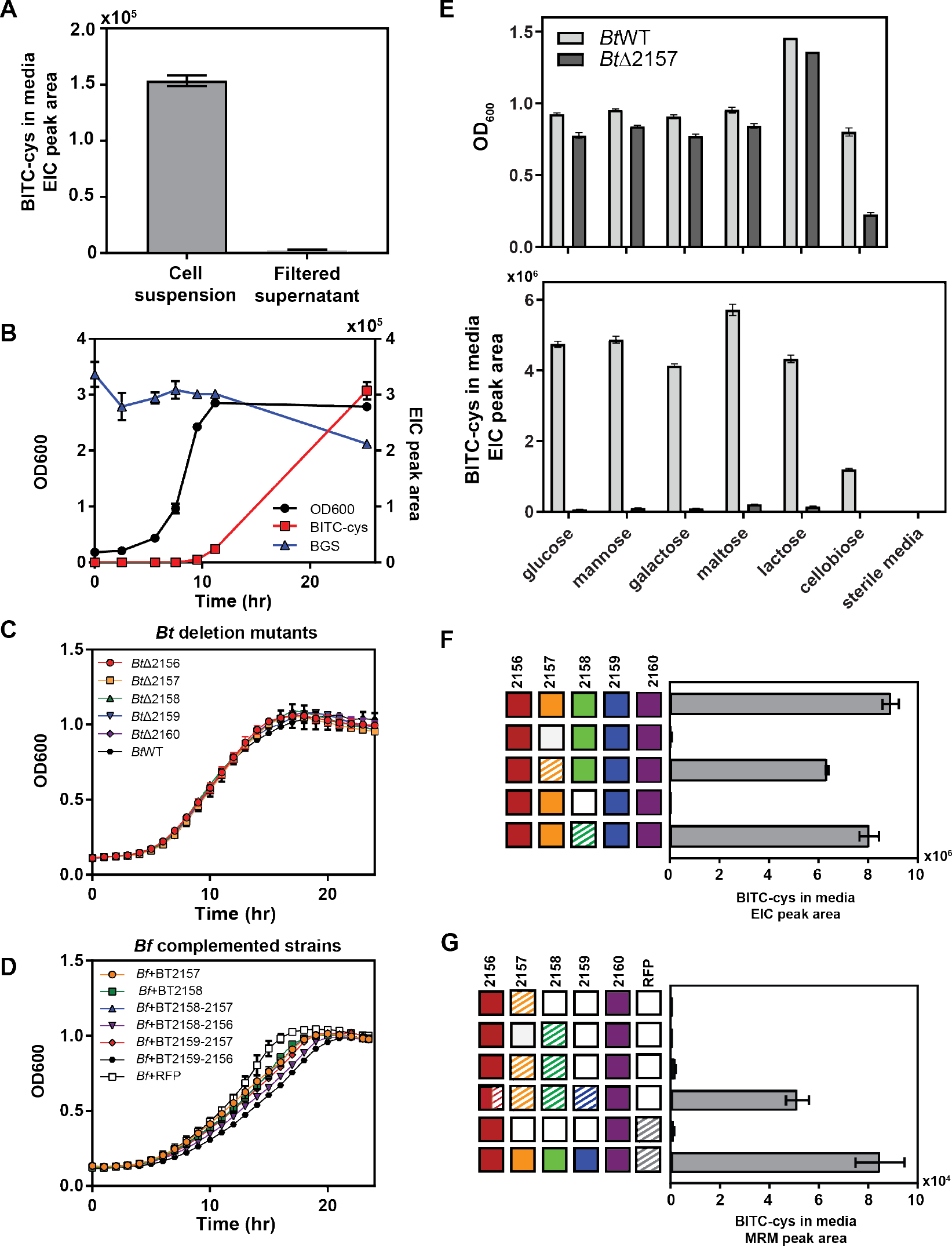
Growth and activity of *Bt* and *Bf* strains in rich media. (A) BITC-cys production 24 hours after addition of BGS to 1) wild-type *Bt* cultured for 24 hours in TYG media, and 2) filtered cell-free spent supernatant from wild-type *Bt* cultured for 24 hours in TYG media. Data shown is the mean ± SD of three biological replicates. Extracted ion count (*m/z* 271.0570) with LC-MS was used to track quantities of protonated BITC-cys. (B) Time courses of growth and glucosinolate conversion for wild-type *Bt* in TYG media supplemented with BGS, measured by optical density (OD) at 600 nm (black), BITC-cys production in the media (red), and BGS depletion (blue). BITC-cys and BGS were monitored by LC-MS and quantified using EIC (BITC-cys: *m/z* 271.0570; BGS: *m/z* 168.0478, corresponding to the -SO4-glucose in-source fragment). Data shown is the mean ± SD of three biological replicates. (C) Growth curves of *Bt* single deletion mutants in TYG media with BGS, measured by OD at 600 nm. Data shown is the mean of three biological replicates ± SD. (D) Growth curves of *Bf* strains expressing subsets of BT2159-BT2156 in TYG media with BGS, measured by OD at 600 nm. Data shown is the mean ± SD of three biological replicates. (E) Growth (top) and BITC-cys production (bottom) of wild-type *Bt* (light grey bars) and *Bt*Δ2157 (dark grey bars) in TY media supplemented with different disaccharides and monosaccharides (0.5% w/v) and 0.5 mM BGS after 24 hours. Data shown is the mean ± SD of three biological replicates. (F) BITC-cys produced from BGS by *Bt*Δ2157 and *Bt*Δ2158 mutants complemented with with BT2157 and BT2158, respectively, grown in TYG media for 24 hours. Each row in the grid of boxes represents the genes expressed in each complemented strain; cross-hatched boxes represent constitutive extra-chromosomal expression from plasmid, while solid boxes represent natively expressed genes. Data shown is mean ± SD of three biological replicates. (G) BITC-cys produced from BGS by *Bt*Δ2159-57 mutant complemented with different sets of the operon proteins. Cross-hatched boxes represent constitutive extra-chromosomal expression from plasmid, while solid boxes represent natively expressed genes. Half solid-half cross hatched boxes represent genes that are both expressed natively and extra-chromosomally. Data shown is mean ± SD of three biological replicates.

**Figure S2.**
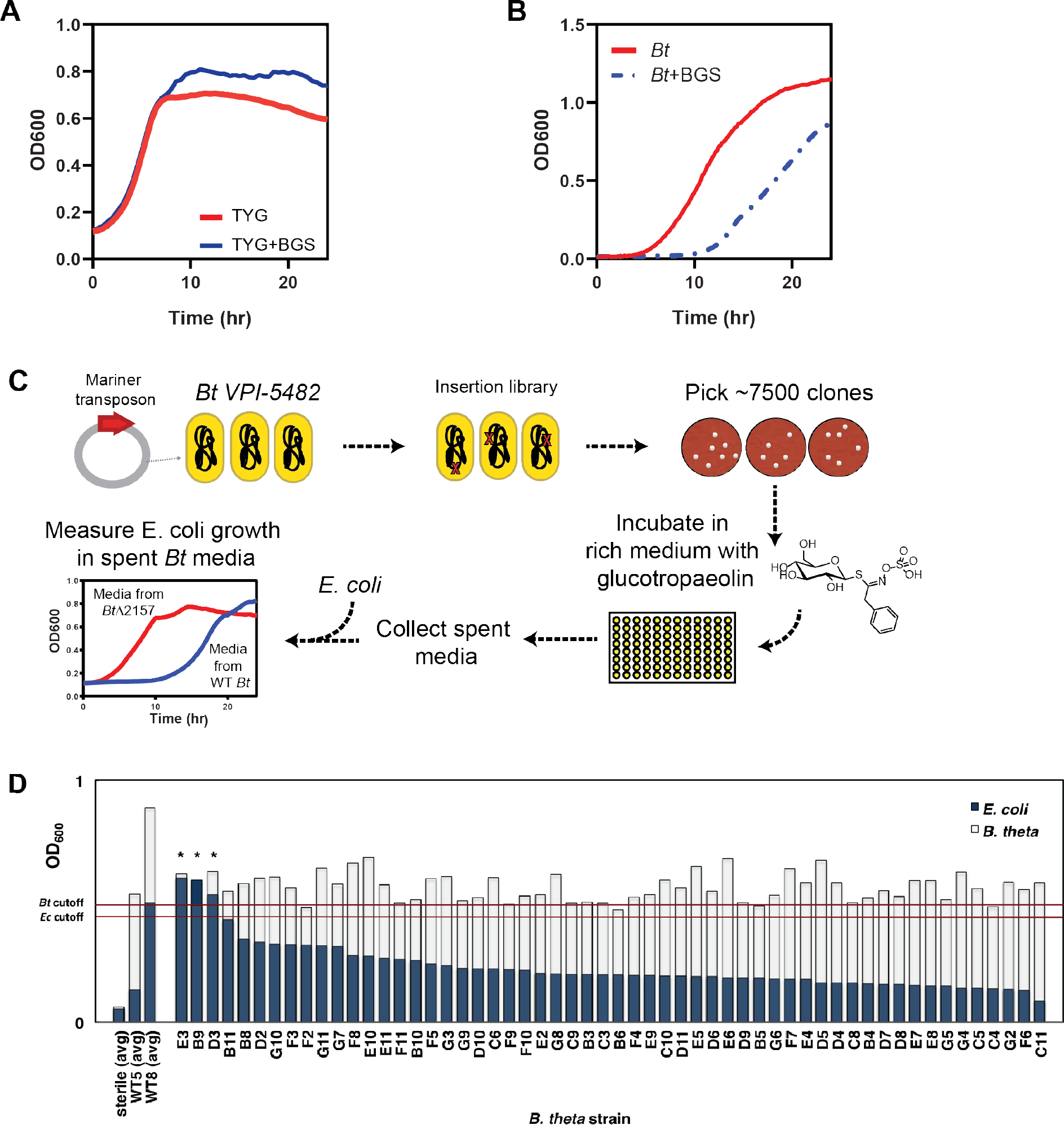
High throughput screen to identify the genes necessary for glucosinolate metabolism. (A) Growth of *Bt* in TYG media (red) and TYG media supplemented with glucotropaeolin (blue). *Bt* growth was not inhibited by glucosinolate conversion to isothiocyanate. Data shown is the average of three biological replicates. (B) Growth of *E. coli* in media spent by wild-type *Bt* cultured for 24 hours in TYG media (red solid line) or TYG media supplemented with glucotropaeolin (blue dashed line). *E. coli* growth was inhibited by glucosinolate conversion to isothiocyanate in *Bt* spent media. (C) Schematic of the high throughput screen developed to identify the genes involved in glucosinolate metabolism, using a transposon insertion library in *Bt* VPI-5482. About 7500 members of the insertion library were screened for loss of glucosinolate metabolizing activity. The bottom left figure shows growth curves for *E. coli* grown in media spent by a *Bt* mutant deficient in glucosinolate metabolizing activity (*Bt*Δ2157, red line) relative to *E. coli* grown in media spent by a *Bt* strain with glucosinolate metabolizing activity (wild type *Bt*, blue line). (D) Representative results of one microplate from high-throughput screen of *Bt* mutant library. Light gray bars in foreground represent the terminal optical densities of *Bt* grown to stationary phase (25 hours). Dark blue bars in background represent the optical density of *E. coli* after 8 hours of growth in spent *Bt* media. Averaged optical densities of sterile and wild-type controls of *Bt* VPI-5482 (WT5) and *Bt* VPI-8736 (WT8), a strain identified with limited glucose metabolizing activity, are shown as the left-most three bars. Mutants with reduced glucosinolate metabolizing activity (marked with *) were identified based on the following two criteria: first, uncompromised growth relative to WT5 (*Bt* mutant OD ≥ average – 1 standard deviation of WT5 OD, “*Bt* cutoff”) and second, loss of the ability to inhibit subsequent *E. coli* growth (*E. coli* OD ≥ average – 1 standard deviation of *E. coli* OD grown in the supernatant of WT8, “*Ec* cutoff”). Approximately 2.9% (218 mutants) of the top hits from the high-throughput screen were chosen for rescreening based on these criteria. Fifty nine mutants were successfully sequenced based on the consistency of their performance in a second and third round of rescreening, using the same criteria employed for the primary screen.

**Figure S3.**
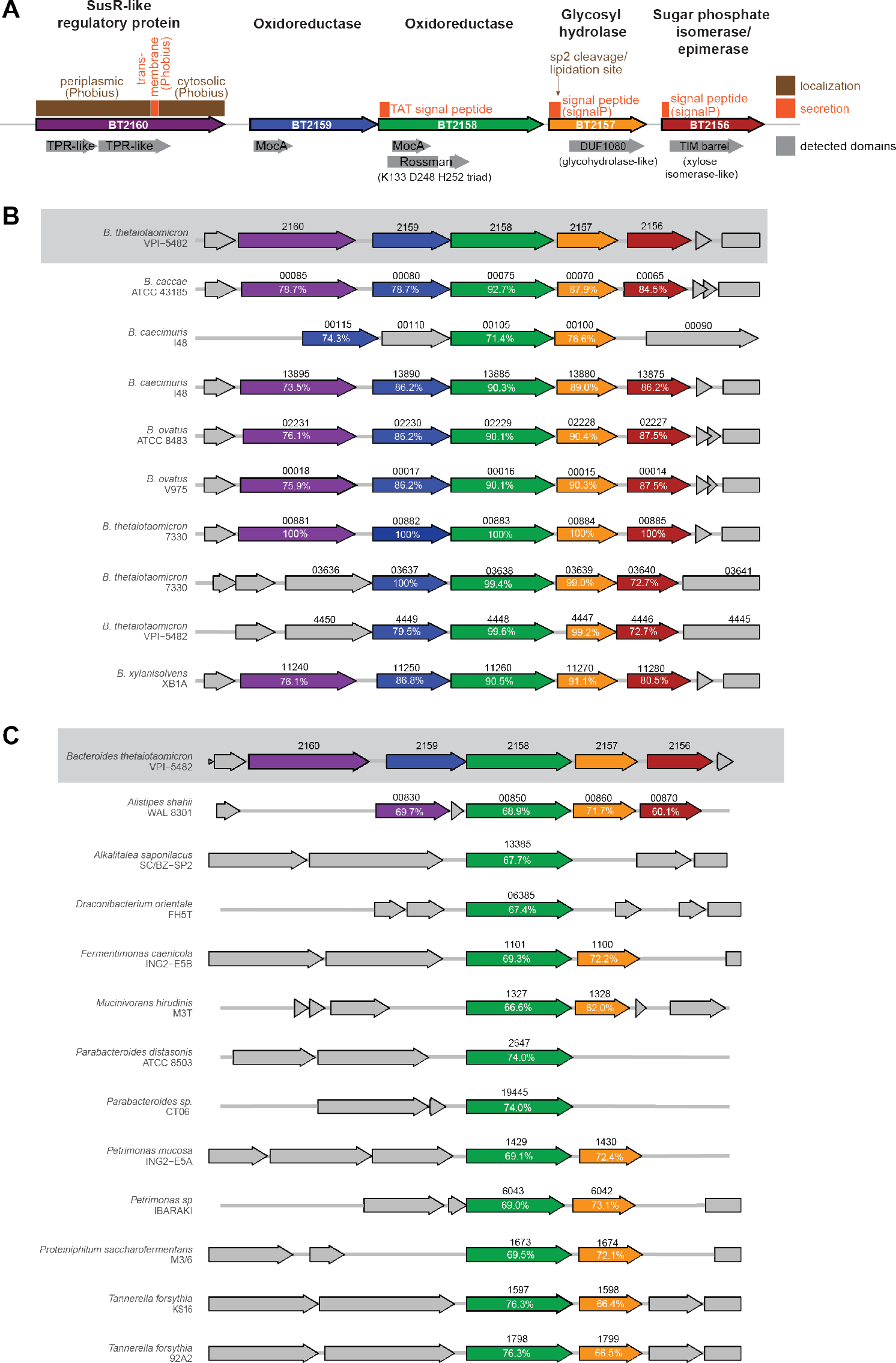
Conservation of the BT2160-2156 operon across bacterial strains. (A) Annotations and predicted domains in the BT2160-BT2156 operon. Detected domains in the operon sequence are shown below the gene arrows. Putative annotations, predicted localization and secretion domains are shown above. (B) Homologs of BT2160-2156 conserved across various *Bacteroides* strains. Homologs were identified using a BLAST search for genes with greater than 60% amino acid similarity and greater than 60% coverage relative to the *Bt* VPI-5482 gene. Percent identity on the amino acid level is shown in white for each gene analyzed. Within *Bt* VPI-5482, BT4449-4446 are 69.5% to 99.6% identical at the amino acid sequence level; however, none of the genes encoding these proteins were identified in the transposon screen and do not appear to be necessary for GS metabolism to ITC. (C) Homologs of BT2160-2156 identified in non-*Bacteroides* genera

**Figure S4.**
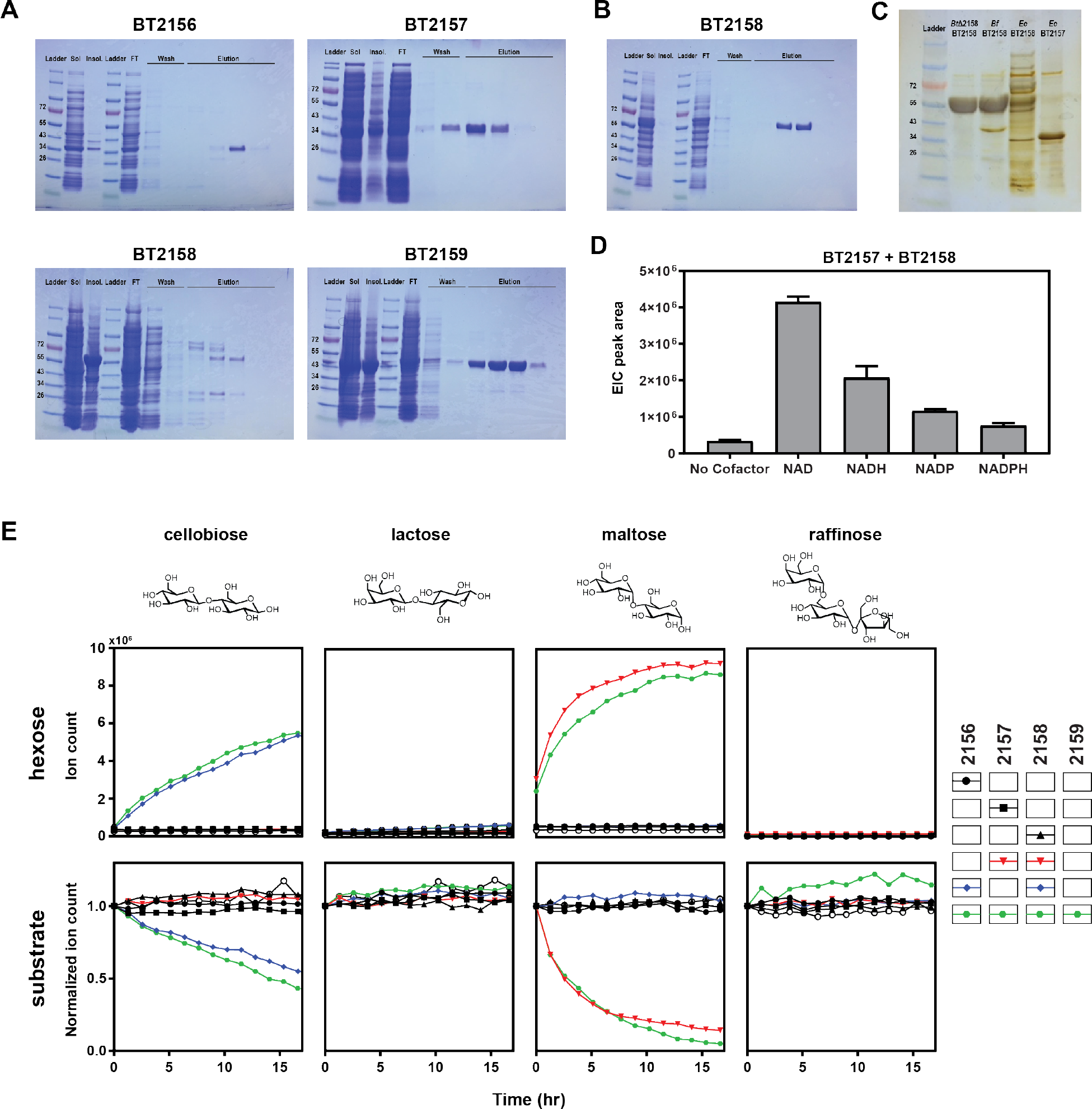
*In vitro* reactions with purified recombinant *Bt* proteins. (A) Coomassie stained SDS-PAGE gels for the Ni-NTA purification of His-tagged BT2159-BT2156 proteins recombinantly expressed in *E. coli* BL21 DE3. Ladder markings show protein sizes in kDa. Sol: total soluble fraction; insol: total insoluble fraction, FT: column flow-through. Total culture volumes were 2 L for BT2156, BT2157, and BT2159. Culture volume was 6 L for BT2158. Less than 1 mg of BT2158 was purified per liter of *E. coli* culture. Despite attempts to optimize expression, most of the BT2158 that was produced remained in the insoluble fraction of the *E.coli* lysate. (B) Coomassie stained SDS-PAGE gel for the Ni-NTA purification of His-tagged BT2158 expressed in *Bt*Δ2158, with a yield of ~10 mg of purified protein per liter of *Bt* culture. The total culture volume was 1 L. (C) Silver stained SDS-PAGE gel of purified fractions of BT2158 expressed in *Bt*Δ2158, *Bf*, and *E. coli*. Purified BT2157 expressed in *E. coli* provided as a reference. (D) *In vitro* conversion of BGS by the combination of purified BT2157 and BT2158 in the presence of 1 mM different nicotinamide cofactors, quantified using EIC peak area of BITC-cys (*m/z* 271.0570), detected by LC-MS. Data shown is the mean ± SD of two replicates. (E) Proteins expressed in *E. coli* (BT2156, BT2157, BT2159) or *Bt*Δ2158 (BT2158) were purified and combined in N-methyl morpholine (NMM) buffer with NAD and di- or trisaccharide substrate. Integrated extracted ion count peaks areas corresponding to hexose release (top row; *m/z* 179.0561) and substrate consumption, normalized against initial substrate ion count, (bottom row; *m/z* 341.1089 for disaccharides and *m/z* 503.1618 for raffinose) were measured. Data shown is one replicate, representative of triplicate experiments.

**Figure S5.**
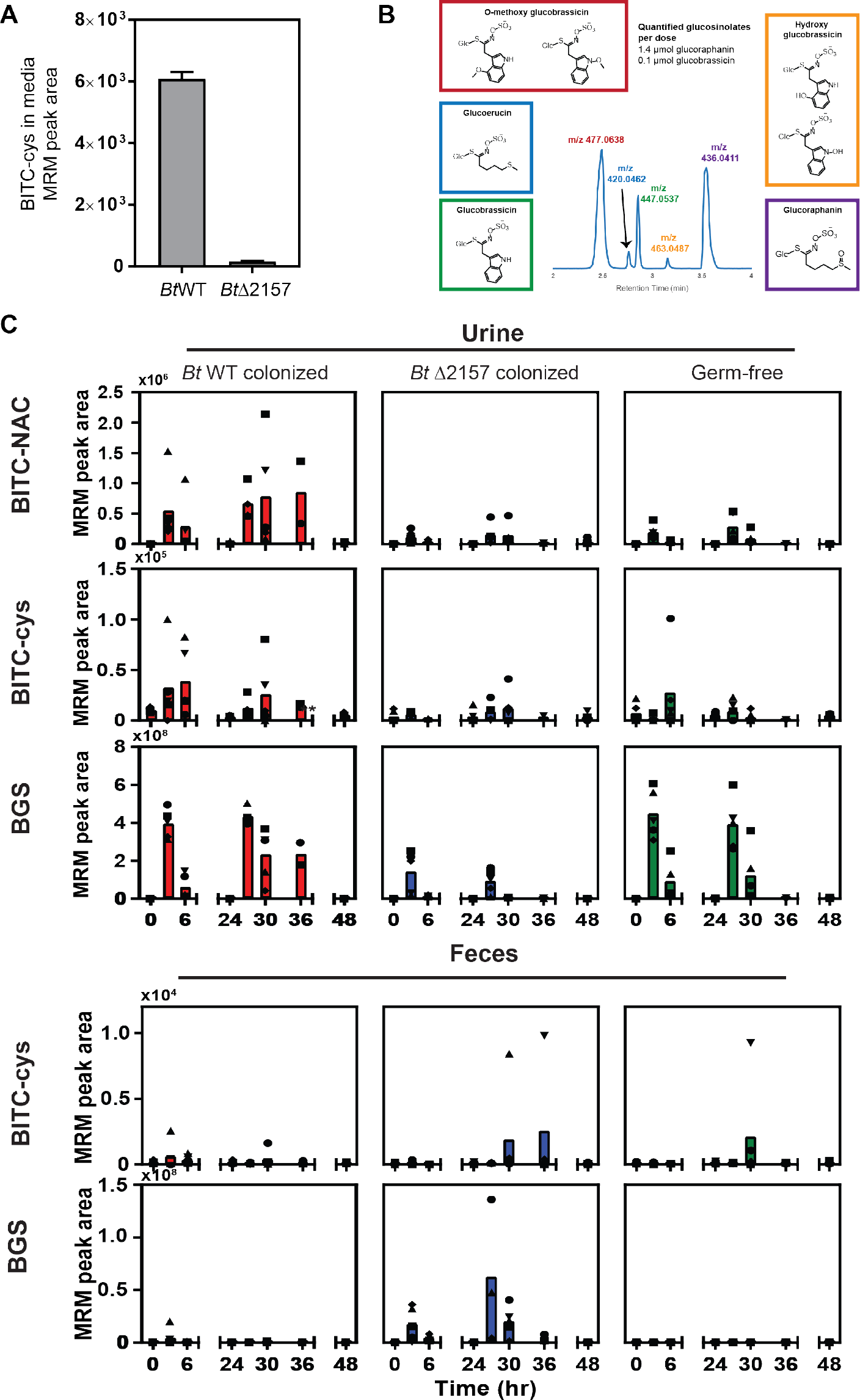
Monocolonization of gnotobiotic mice with mutant *Bt*. (A) Feces from mice colonized with wild-type *Bt* (light grey bar) or *Bt*Δ2157 colonized mice (dark grey bar) were collected prior to glucosinolate dosing, plated at limiting dilutions, and single colonies were cultured in TYG medium with BGS. Multiple reaction monitoring (MRM) by LC-MS/MS was used to track the transition of the protonated BITC-cys molecule with *m/z* 271.0 to a product ion with *m/z* 122.0. Data shown are mean ± SD of five biological replicates. (B) Annotated representative extracted ion chromatogram of glucosinolates identified in broccoli extract. Glucosinolates corresponding to the extracted exact masses are shown in boxes. (C) BITC-NAC and BITC-cys mercapturic acid conjugates and unconverted BGS excreted in urine and feces after doses of pure glucotropaeolin (top), and BITC-cys and unconverted BGS in feces collected after doses of pure glucotropaeolin (bottom). BITC-cys in feces was the product of *in situ* conjugation of free BITC with exogenously applied cysteine. Bars represent the mean values for each group, with individual data overlaid (different shapes correspond to individual mice). Metabolites were measured by LC-MS/MS and quantified by MRM (BITC-NAC: *m/z* 313.1→ *m/z* 164; BITC-cys: *m/z* 271.1→ *m/z* 122.0; BGS: *m/z* 408.0→ *m/z* 96.9). Mice mono-colonized with mutant *Bt*Δ2157 compared to wild-type *Bt* showed increased glucosinolate concentration in fecal samples (**Figure S6B**), which may be indicative of indirect impacts of glucosinolate conversion, other physiological activities encoded by this operon (Liu et al., 2019), or effects specific to monocolonized mouse models.

**Table S1.**
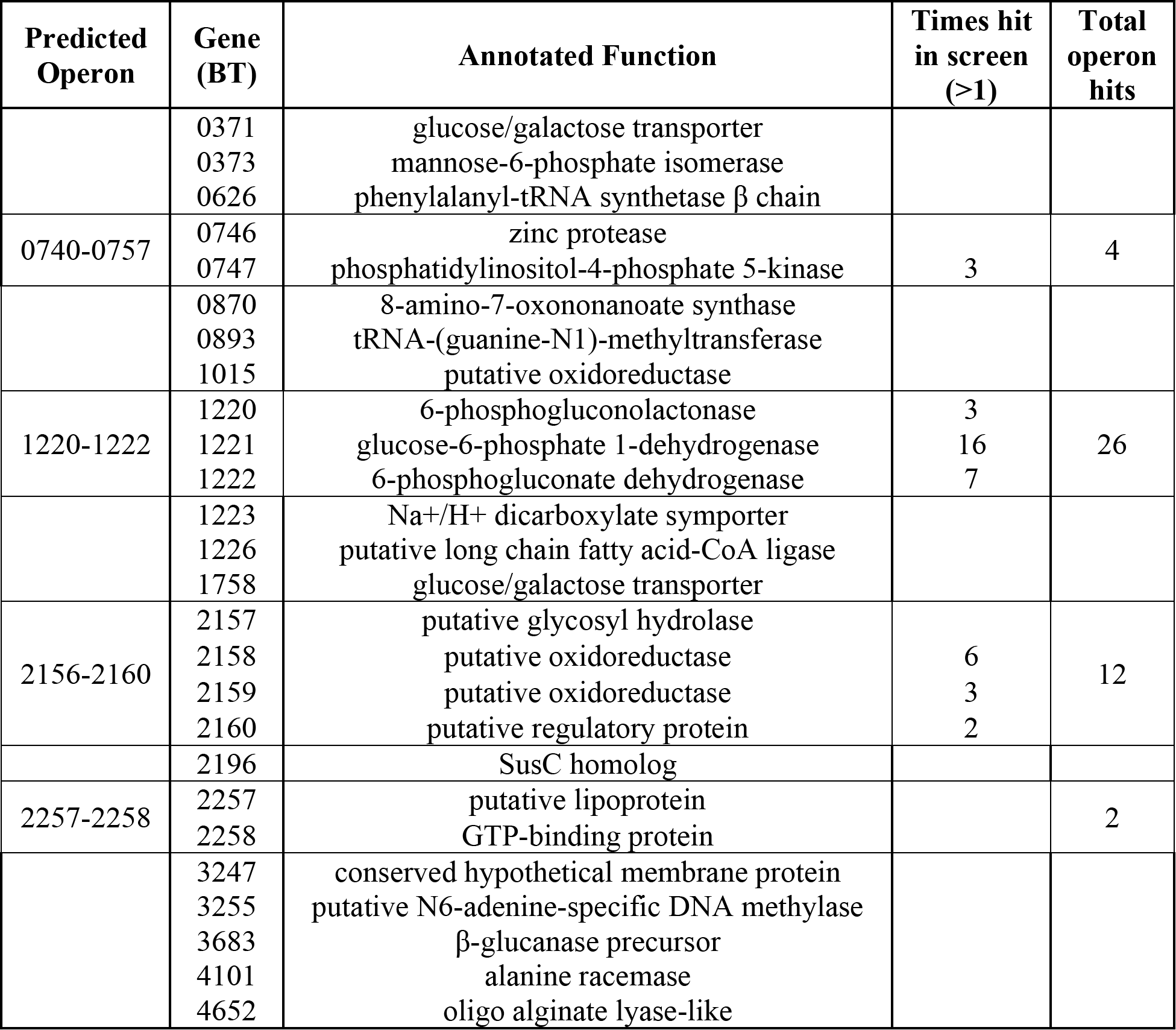
Genes identified in the genome wide transposon screen.

## METHODS

### *Bacteroides* culturing conditions

*Bacteroides* strains were streaked from glycerol stocks on either brain heart infusion agar supplemented with 10% horse blood (BHI-BA) plates or brain heart infusion agar supplemented with 5 μg/mL hemin and 0.5 μg/mL menadione (BHIS) plates as indicated. Liquid cultures were grown in tryptone-yeast extract medium, prepared as described previously (Whitaker et al., 2017) and freshly supplemented with 0.5 mg/mL cysteine as a reducing agent and 0.5% glucose (TYG). All *Bacteroides* culturing was performed under anaerobic conditions, using the GasPak anaerobic system (BD) or an anaerobic chamber (Coy, 10% CO_2_, 5% H_2_).

### Transposon mutagenesis of *Bacteroides thetaiotaomicron* VPI-5482

The transposon library was constructed using the Mariner transposon vector pSAM_Bt to mutagenize *Bacteroides thetaiotaomicron (Bt)* VPI-5482 (Goodman et al., 2009). *Bt* VPI-5482 was inoculated from glycerol stocks into TYG and cultured anaerobically at 37°C for 16-18 hours. *Escherichia coli (E. coli)* harboring pSAM_Bt was similarly inoculated into LB with 50 μg/mL ampicillin and grown aerobically with shaking at 37°C for 16-18 hours. *Bt* and *E. coli* were subcultured 1:20 into TYG and 1:100 into LB with ampicillin, respectively, and grown to exponential phase as determined by OD_600_ values between 0.4 and 0.6. Cells were pelleted by centrifugation at 17,000 *g* for 20 minutes. For the conjugation reaction, the *Bt* and *E. coli* pellets were resuspended in 1 mL of TYG, combined, plated as three 100μL spots on BHI-BA, and then incubated aerobically for 8 hours at 37°C. Conjugation reactions were pooled and resuspended in phosphate buffered saline (PBS). Pooled resuspensions were then plated as 100 μL aliquots onto each of 50 BHI-BA plates supplemented with 25 μg/mL erythromycin and 200 μg/mL gentamicin and grown anaerobically at 37°C for 20-30 hours. Following selection, single colonies were inoculated into 200 μL TYG containing 25 μg/mL erythromycin and 200 μg/mL gentamicin and grown anaerobically at 37°C for 48 hours. Glycerol stocks of individual mutants were prepared by supplementing the liquid cultures with an equal volume of 60% glycerol in TYG (30% glycerol final concentration) and stored at −80°C until screening for glucosinolate metabolizing activity.

### High-throughput bioassay for loss of glucosinolate metabolizing activity

To screen for disruption in candidate glucosinolate metabolizing genes in the *Bt* transposon insertion library, the isothiocyanate (ITC)-sensitive *E. coli* strain TOP10 harboring the irrelevant plasmid pENTR-sGFP (Life Technologies), included to permit growth on kanamycin-containing media, was inoculated from a glycerol stock into LB containing 50 μg/mL kanamycin and incubated aerobically with shaking at 37°C for 16-18 hours. This *E. coli* culture was diluted 1:10 into LB with kanamycin in preparation for use as the indicator strain for the bioassay (described below). In parallel, *Bt* transposon mutants were inoculated from glycerol stocks into TYG in microwell plates and incubated anaerobically at 37°C for 22-30 hours. The *Bt* mutants were subcultured 1:50 into TYG supplemented with 0.5 mM glucotropaeolin (BGS) and incubated anaerobically at 37°C for an additional 23-25 hours. The endpoint OD_600_ values of the *Bt* cultures were measured, and then the cells were pelleted by centrifugation at 5,000 *g* for 15 minutes. Next, 90 μL of supernatant was transferred to microwell plates containing 10 μL of the diluted *E. coli* culture described above (1:100 overall *E. coli* dilution). Microplates were incubated aerobically with agitation at 37°C. Growth of *E. coli* was monitored by measuring OD_600_ at the following timepoints: 0 h, 3 h, 6 h, 9 h, and 16 h.

Wild-type *Bt* VPI-5482 and *Bt* 8736 were included on each microplate as positive and negative control strains, respectively, for glucosinolate metabolizing activity. Approximately 2% of mutants (152 mutants) from the high-throughput screen were chosen for rescreening based on the following two criteria: 1) uncompromised *Bt* growth relative to wild-type *Bt* VPI-5482 (*Bt* mutant OD_600_ ≥ average – 1 standard deviation of *Bt* VPI-5482 OD_600_), and 2) loss of ability to inhibit subsequent *E. coli* growth (*E. coli* OD_600_ ≥ average – 1 standard deviation of *E. coli* OD_600_ grown in supernatant spent by *Bt* 8736). Of this top 2% of mutants, 77 were chosen for sequencing based on the consistency of their performance in a second and third round of rescreening, using the same criteria employed for the primary screen.

The following instrumentation was used at the High-Throughput Bioscience Center at the Stanford School of Medicine: Infinite M1000 plate reader (Tecan) to measure OD_600_, Matrix Wellmate (Thermo Scientific) to dispense media, and Bravo Liquid Handler (Agilent) to transfer spent media.

### Identification of Transposon Mutation Sites

Transposon insertion sites in *Bt* VPI-5482 mutants were identified using a semi-random PCR approach, adapted from Goodman et al. (Goodman et al., 2009). Genomic regions adjacent to transposon insertion sites were amplified by PCR using the transposon-specific primer 5’-ACGTACTCATGGTTCATCCCGATA-3’ and one of three degenerate primers 5’-GGCCACGCGTCGACTAGTACNNNNNNNNNNGATGC-3’, 5’-GGCCACGCGTCGACTAGTACNNNNNNNNNNGGCCG-3’, 5’-GGCCACGCGTCGACTAGTACNNNNNNNNNNGTAAT-3’, where N represents either A, T, G, or C synthesized using the hand-mix option from Integrated DNA Technologies. Products from the first PCR were purified (QIAquick kit, Qiagen) and used as template for a second round of PCR, using the nested transposon specific primer 5’-GCGTATCGGTCTGTATATCAGCAA-3’ and primer 5’-GGCCACGCGTCGACTAGTAC-3’. Following agarose gel electrophoresis, amplicons were purified from an agarose gel (Zymoclean kit, Zymo Research) and submitted for Sanger sequencing using the nested transposon-specific primer 5’-TCTATTCTCATCTTTCTGAGTCCAC-3’. Loci corresponding to the genomic regions adjacent to transposon insertions were identified using BLAST.

In this manner, transposon insertion sites were identified for 65 *Bt* mutants that displayed a loss of glucosinolate metabolizing activity. More than two thirds of these mutations occurred in two putative operons. One operon (BT1220-1222) encodes genes annotated as the pentose phosphate pathway and was deemed likely to eliminate glucosinolate metabolizing activity via pleiotropic effects. The other operon (BT2156-2160) carried annotations suggestive of carbohydrate metabolizing activity was selected for further molecular genetic and biochemical characterization.

### Targeted mutagenesis of *Bt* VPI-5482

All targeted deletion mutants in *Bt* were constructed in a 5-fluoro-2’deoxyuridine (FUdR) resistant *Bt* VPI-5482 mutant lacking thymidine kinase (*Bt*Δ*tdk*) using counterselectable allelic exchange (Koropatkin et al., 2008). Genomic sequences spanning 1.5 kb upstream and downstream of the gene to be deleted were amplified from *Bt* VPI-5482 genomic DNA by PCR. Amplicons were then assembled using fusion PCR into an in-frame start-stop codon pair flanked by the ~1.5 kb genomic regions surrounding the target gene. Purified inserts were cloned into the pExchange-tdk suicide vector digested with PstI and NotI using the Gibson assembly method (Gibson et al., 2009)and transformed into *E. coli* S17-1 λpir chemically competent cells. Plasmids from several transformants were miniprepped and sequenced to confirm correct assembly, and then conjugated into *Bt*Δ*tdk*. For conjugation, *E. coli* S17-1 λpir harboring the gene-targeting pExchange shuttle vectors were grown on LB plates containing 100 μg/mL ampicillin and inoculated into LB containing 100 μg/mL ampicillin and grown for 17-20 hours at 37°C with agitation. In parallel, *BtΔtdk* was grown anaerobically on BHI-BA plates, and then inoculated into TYG and grown anaerobically for 17-20 hours at 37°C. Cells were pelleted by centrifugation at 1400 *g* for 15 minutes and washed with LB. Pelleted strains containing the donor plasmid were resuspended in LB and then used to resuspend *BtΔtdk*. Mating mixtures were spotted onto BHI-BA plates and incubated aerobically at 37°C for 24 hours. To select for the first round of recombination, mating cultures were streaked onto BHI-BA plates containing 200 μg/mL gentamycin and 5 μg/mL erythromycin and grown anaerobically for 48 hours. To counter-select for the second recombination event, colonies from the first round were streaked onto BHI-BA plates containing 200 μg/mL FUdR and grown for 48-72 hours. PCR amplification of genomic DNA flanking the gene was used to confirm deletion, using the following sets of primers:

BT2159_F: 5’-GCTAGTTTTGCGATATCAGTTTTCG-3’
BT2159_R: 5’-GGCAACTTCCATCCTTCACG-3’
BT2158_F: 5’-AGCTAATGGATAAAATACTTTCTTCATAATAATAACTATTTAATTTTTCT-3’
BT2158_R: 5’-ATCCGTGCTGCAGCCAATATAAACGAATTTGTGCCA-3’
BT2157_F: 5’-AACAAGACACTGGAATGGGAC-3’
BT2157_R: 5’-CACCGCGGTGGCGGCCGCTCTAGCAGTATAGCTGTCGATTAGTATG-3’
BT2156_F: 5’-CGATAAGCTTGATATCGAATTCCTGCAGCAGTATAGCTGTCGATTAGTATG-3’
BT2156_R: 5’-ATCCGTGGATCCCACGATAAAAGTAAGTTAAGA-3’

PCR amplification of the gene was used to confirm that the gene had not been inserted into another region of the genome using the following sets of primers:

BT2159_F: 5’-TGCGATACAGATCCTACCACGC-3’
BT2159_R: 5’-CAATGTGAAGAGCCCGACAACC-3’
BT2158_F: 5’-CGAAACAATTTGCAGCCGAAC-3’
BT2158_R: 5’-GGCAACTTCCATCCTTCACG-3’
BT2157_F: 5’-TGCAAGCCAGCAAATTCAGC-3’
BT2157_R: 5’-CAGTCCAGAACTTTCACGCG-3’
BT2156_F: 5’-CTGCCGGGCTGAAGGTTTTATC-3’
BT2156_R: 5’-TCAGCAATACACTGGTCCCACC-3’

### Cloning and heterologous expression of BT2159-BT2156 in *Bacteroides fragilis*

Expression of subsets of BT2159-BT2156 in the heterologous host *Bacteroides fragilis* (*B. fragilis*) was achieved using the extra-chromosomal *Bacteroides* expression vector, pFD340 (Smith et al., 1992). Target gene sequences, including the ~50 bp region upstream of the gene to encapsulate the native ribosome binding site, were amplified from *Bt* VPI-5482 genomic DNA by PCR. Amplicons encoding sequences for BT2159-BT2157 and BT2158-BT2156 were purified, and then inserted into pFD340 plasmid digested with BamHI using the Gibson assembly method. All other single and multi-gene amplicons were purified, digested with SacI and BamHI, and then ligated with T4 DNA ligase into pFD340 plasmid digested with the same enzymes. An empty vector control was generated by amplifying a sequence encoding RFP, digesting the purified amplicons with BamHI and PstI, and then ligating it using T4 DNA ligase into pFD340 digested with the same enzymes. Assembled vectors were transformed into chemically competent *E. coli* TOP10 and conjugated into *B. fragilis* via triparental mating, similarly to the procedure described above, but with the inclusion of helper strain *E. coli* RK231 in the mating mix (Smith et al., 1992). Selection was performed on BHI-BA plates containing 200 μg/mL gentamycin and 5 μg/mL erythromycin.

### Bacterial assays for glucosinolate metabolism

*Bacteroides* strains were streaked from glycerol stocks onto BHIS plates and incubated anaerobically at 37°C for 22-26 hours. For strains harboring the pFD340 plasmid, 5 μg/mL of erythromicin and 200 μg/mL of gentamicin were included. Single colonies were inoculated into 200 μL of TYG, containing 5 μg/mL erythromycin if harboring a pFD340 plasmid, and grown anaerobically at 37°C without shaking. After 17-24 hours of growth, strains were subcultured into 200 μL fresh TYG supplemented with 0.5 mM glucotropaeolin (BGS) and 5 μg/mL erythromycin, if required for plasmid selection. Following anaerobic incubation at 37°C for 24 hours, cells were pelleted by centrifugation at 1400 *g* for 20 minutes. The supernatant was diluted 1:10 with [10% (v/v) acetonitrile and 90% (v/v) water with 0.1% (v/v) formic acid], filtered through 0.45 μm PTFE filters, and analyzed by either liquid chromatography-mass spectrometry (LC-MS) or liquid chromatography-tandem mass spectrometry (LC-MS/MS), as described below.

### *Bacteroides* growth curves and ITC time course

For growth curves of *Bt* and *Bf* mutants, strains were cultured using the same method as described for glucosinolate metabolism assays above. Following subculturing in TYG supplemented in 0.5 mM BGS, cultures were transferred into an anaerobic chamber and OD_600_ measurements were taken every 10 minutes using an Epoch 2 plate reader (Biotek) held at 37°C.

For simultaneous measurements of growth and ITC production over time, 12 mL of wild-type *Bt* was cultured in the presence of 0.5 mM BGS as previously described for glucosinolate metabolism assays, in an anaerobic chamber at 37°C. Growth of *Bt* cultures was measured over a course of 24 hours by sampling 700 μL of culture and measuring OD_600_ using a Genesys 20 spectrophotometer (Thermo Scientific). After OD_600_ measurement at each sampled time point, 500 μL of culture was frozen in liquid nitrogen and stored at −80°C. After completion of the time course, frozen samples were thawed and centrifuged at 13,000 *g* for 10 minutes, and spent media was diluted and analyzed by LC-MS as described above.

### Recombinant protein expression and purification

Sequences encoding BT2156, BT2157, and BT2159 were amplified from *Bt* VPI-5482 genomic DNA by PCR. Purified amplicons for BT2156, BT2157, and BT2159 were digested with BamHI and XhoI and ligated using T4 DNA ligase into the C-terminal His-tagged expression vector pET-24b digested with the same enzymes. Plasmids were transformed into *E. coli* TOP10 chemically competent cells, selected, miniprepped, and verified by Sanger sequencing (ELIM Biopharm). For protein expression, purified plasmid was subsequently transformed into *E. coli* BL21 DE3 chemically competent cells.

*E. coli* BL231 DE3 strains were grown for 18 hours at 37°C from glycerol stocks streaked on LB agar plates containing 50 μg/mL kanamycin. Single colonies were inoculated into 20 mL of LB with 50 μg/mL kanamycin. Following growth at 37°C for 18 hours, strains were subcultured into 2L of LB with kanamycin and grown at 37°C with agitation until an OD_600_ of 0.6 was reached. Cultures were induced with 0.1 mM ispropyl β-D-1 thiogalactopyranoside (IPTG), after which they were incubated at 30°C for 6 hours. Cells were pelleted by centrifugation at 5500 *g* for 5 mins, and supernatant was discarded. Cell pellets were then frozen in liquid nitrogen and stored at −80°C.

For expression of His-tagged BT2158 in *Bt* VPI-5482, the gene was amplified from genomic DNA, including 50 bp upstream of the start codon to encapsulate the native RBS, as well as an appended sequence encoding a C-terminal 6xHis tag. Purified amplicon was digested with BamHI and XhoI and ligated using T4 DNA ligase into pFD340 digested with the same enzymes. To achieve higher levels of protein expression, the IS4351 promoter sequence in the pFD340 vector was replaced with the phage promoter P_BfP1E6 sequence (Whitaker et al., 2017) using the PstI and BamHI restriction sites. Ligation products were transformed into *E. coli* TOP10 chemically competent cells and miniprepped plasmids were confirmed using Sanger sequencing. The construct was expressed in single deletion mutant *Bt*Δ2158 via triparental conjugation with helper strain RK231, as described above. The transformed strain was streaked onto BHIS plates containing 200 μg/mL gentamycin and 5 μg/mL erythromycin and incubated at 37°C for 48 hours under anaerobic conditions. Single colonies were inoculated into 24 mL of TYG containing 5 μg/mL erythromycin and grown at 37°C for 24 hours. The strain was subcultured into 1 L of TYG containing the same antibiotics and incubated anaerobically at 37°C for another 20 hours. Cells were pelleted by centrifugation at 6000 *g* for 5 minutes, and supernatant was discarded. Cell pellets were frozen in liquid nitrogen and stored at −80°C.

For protein purification, *E. coli* and *Bt*Δ2158 cell pellets were thawed by resuspension in 25 mL of lysis buffer (50 mM potassium phosphate, 300 mM NaCl, 10 mM imidazole, pH 7.8) and lysed with six 20 second bursts of sonication at 60% amplitude. Cell debris was pelleted by centrifugation at 38,000 *g* for 30 minutes at 4°C and discarded. All subsequent processes were performed at 4°C. Lysate was equilibrated with Ni-NTA agarose resin (Thermo Scientific) for 1 hour with rocking and loaded onto a 1.5×10 cm Econo-column chromatography column (Bio-rad). The resin was washed with 20 mL lysis buffer and 20 mL wash buffer (50 mM potassium phosphate, 300 mM NaCl, 20 mM imidazole, pH 7.8), followed by elution with successively higher concentrations of imidazole (50 mM potassium phosphate, 300 mM NaCl, 50/100/200/500 mM imidazole, pH 7.8). Elution fractions were analyzed by SDS-PAGE and the fractions with the desired protein were combined. Pooled fractions were buffer exchanged into storage buffer (50 mM potassium phosphate, pH 7.8) and concentrated using Amicon Ultra-4 centrifugal filter units with 10 kDa molecular weight cut off (Millipore). Glycerol was added to a final concentration of 10% (v/v), after which fractions were aliquoted, frozen in liquid nitrogen, and stored at −80°C. Total protein concentration was quantified using absorbance at 280 nm, measured using a Nanodrop 1000 spectrophotometer (Thermo Scientific).

### *In vitro* endpoint assays with purified protein

Standard *in vitro* endpoint protein reactions consisted of 0.5 μL of 100 mM BGS substrate, 1 μL of 50 mM NAD_+_, 200 pmol of each protein, and reaction buffer (10 mM potassium phosphate buffer, pH 7.2) up to a total volume of 50 μL. Each reaction contained final concentrations of 1 mM each of BGS and NAD_+_, and 4 pmol/μL of each protein. Reactions were initiated by the addition of substrate and incubated at 37°C for 15 hours, prior to quenching with 90 μL acetonitrile and 10 μL 15 mg/mL cysteine, to promote conjugation of free ITC and enable detection of ITC-cysteine conjugates by MS. Quenched reactions were centrifuged at 15,000 *g* for 5 minutes to pellet denatured protein. Supernatant was filtered through a 0.45 μm PTFE filter prior to analysis by LC-MS, as described below.

### *In vitro* kinetic assays with purified protein

All proteins were buffer exchanged using Amicon ultra-4 centrifugal filters with 10 kDa molecular weight cut off (Millipore) into volatile reaction buffer (10 mM N-methyl morpholinium acetate, pH 7) to ensure MS compatibility. Kinetic reactions monitoring a time course of *in vitro* protein activity consisted of 1 μL of 100 mM substrate, 2 μL of 50 mM NAD^+^, 450 pmol of each protein, and reaction buffer up to a total volume of 100 μL. Each reaction contained final concentrations of 1 mM each of substrate and NAD^+^, and 4.5 pmol/μL of each protein. Reactions were initiated by the addition of substrate, incubated at room temperature in an Agilent 1290 Infinity II multisampler, and sampled hourly for MS analysis for 17 hours, using the conditions described below.

### Mouse study

Animal studies were performed under a protocol approved by the Stanford University Institutional Animal Care and Use Committee. Germ-free Swiss Webster mice (Taconic) were maintained in gnotobiotic isolators under aseptic conditions and fed *ad libitum* with a standard autoclaved chow diet (LabDiet 5K67). Groups of five 10-13 week old male mice were colonized with either wild-type *Bt* or *Bt*Δ2157 by oral gavage of overnight cultures (Marcobal et al., 2011), or maintained germ-free as a negative control. To confirm colonization and maintenance of WT or mutant *Bt* status, fecal pellets were collected six days after gavage. Briefly, 1 μL of fecal pellet was serially diluted, plated on BHI-BA, and grown anaerobically for 24 hours at 37°C. Cultures were inoculated into TYG supplemented with 0.5 mM BGS and grown anaerobically for 24 hours at 37°C. Cells were pelleted, and spent supernatant was assayed for conversion of glucosinolate to isothiocyanate by LC-MS/MS, as described below (Figure S5A). Five days following colonization, the mice were switched to a polysaccharide deficient (PD) diet (Bio-Serv S5805).

Two days following introduction of the PD diet, all mice received two daily doses of 200 μL of 45 mM sterile-filtered BGS by gavage, corresponding to a total dosage of 9 μmol per animal per day. BGS doses were sterile filtered to prevent contamination of the gnotobiotic environment. Urine and fecal samples were collected immediately prior to each treatment, as well as 3 and 6 hours after the first dose and 3, 6, 12, and 24 hours after the second dose. Urine and feces were both stored at −80°C until analysis and analyzed for BGS and BITC content by a combination of LC-MS and LC-MS/MS. Previous studies have shown that the primary metabolic fate of ITCs is conjugation to glutathione, followed by sequential metabolism via the mercapturic acid pathway to ITC-cysteinylglycine and ITC-cysteine (Shapiro et al., 2001). ITC-cysteine conjugates are acetylated to N-acetyl cysteine conjugates, which are then excreted in the urine. Previous studies have established ITC-NAC excretion in urine as a biomarker for crucifer intake and ITC exposure (Hwang and Jeffery, 2003). Urine samples were specifically analyzed for N-acetyl cysteine conjugates of ITC (ITC-NAC) (Figure 1C), as well as ITC-cysteine conjugates, the metabolite upstream of ITC-NAC conjugates in the mercapturic acid pathway. Metabolites from fecal samples were extracted, incubated with an excess of cysteine for reversible thiol conjugation, and analyzed for ITC-cys conjugates by LC-MS/MS.

Five days following the second pure BGS dose, mice received four daily doses of broccoli meals containing broccoli glucosinolates in a food matrix. To maintain gnotobiotic conditions, broccoli meals were comprised of a mixture of autoclaved, glucosinolate-free broccoli slurry and sterile-filtered, concentrated broccoli extract.

Broccoli meals were prepared from broccoli florets purchased from a local grocery store. To prepare concentrated broccoli extract, florets were microwaved for 3 minutes to inactivate native plant myrosinase, frozen in liquid nitrogen, and then lyophilized to dryness. Lyophilized florets were ground using a ball mill homogenizer (Retsch) and then reconstituted in water (13 mL/g dry weight) at 75°C for 20 minutes. Plant material was collected by centrifugation at 1875 *g* for 15 minutes and then discarded. Supernatant was filtered through a 5 μm nylon filter to remove remaining plant material. To concentrate the extract, filtered supernatant, totaling about 485 mL, was frozen in liquid nitrogen, and then lyophilized to dryness. Lyophilized extract was resuspended in 33 mL of deionized water, resulting in about a ~15 fold concentration of the original extract. Concentrated extract was sterile filtered through a 0.2 μm regenerated cellulose filter, aliquoted, frozen in liquid nitrogen, and then stored at −20°C. To quantify glucosinolates in the concentrated extract, 90 μL of extract was diluted into 180 μL of hydrophilic interaction liquid chromatography (HILIC) acetonitrile mobile phase, filtered through a 0.45 μm PTFE filter, and then analyzed by LC-MS, as described below.

For the glucosinolate-free broccoli slurry, raw florets were frozen in liquid nitrogen and then lyophilized to dryness. Lyophilized florets were ground using a ball mill homogenizer, reconstituted in water (8 mL/g dry weight), and then autoclaved at 128°C for 20 minutes, a process that resulted in thermal degradation of glucosinolates. Immediately prior to gavage, 1.8 mL of thawed concentrated extract was added to 1.2 mL of the glucosinolate-free slurry to constitute broccoli meals for a given treatment. To prevent degradation of glucosinolates in the extract, concentrated extracts were stored at −20°C and transferred daily into the gnotobiotic incubators.

All mice received daily doses of 200 μL of broccoli meal by gavage for four consecutive days, corresponding to a quantified dosage of 1.4 μmol of glucoraphanin and 0.13 μmol of glucobrassicin, as well as unquantified amounts of hydroxyglucobrassicin, methoxyglucobrassicin, and glucoerucin. Urine and feces were collected immediately prior to, as well as 3 and 6 hours after the first and third doses. Urine and feces samples were both stored at −80°C until analysis.

Twenty four hours following the final broccoli meal, all mice received one dose of 200 μL of BGS mixed with the glucosinolate-free broccoli slurry, providing a total dose of 4.5 μmol of BGS per mouse. Urine and feces were collected immediately prior to the feeding, as well as 3 and 6 hours following the feeding. Urine and feces were both stored at −80°C until analysis. Mice were sacrificed six hours following the feeding by CO2 asphyxiation in accordance with approved protocols, blood samples were collected in Microtainer SST tubes (BD), and serum was harvested. Serum samples were frozen in liquid nitrogen and stored at −80°C.

Urine samples were thawed and centrifuged at 14,000*g* for 5 minutes. For samples collected following pure BGS treatments, 30 μL of sample was diluted into 30 μL acetonitrile and 240 μL water with 0.1% (v/v) formic acid, and then filtered through a 0.45 μm PTFE filter. For samples in which less than 30 μL of urine was collected, deionized water was added to make up the difference and the dilution was accounted for in corresponding peak area calculations. Filtered samples were analyzed using LC-MS with an Agilent 6520 qTOF, as described below. For urine samples collected from broccoli meal feedings, 15 μL of urine was diluted into 135 μL of HILIC acetonitrile mobile phase. For urine samples with less than 15 μL volume, the quantity of HILIC mobile phase used as diluent was adjusted to maintain a 10-fold dilution factor. Diluted samples were filtered through a 0.45 μm PTFE filter, and then analyzed by LC-MS/MS, as described below.

Fecal pellets were thawed, resuspended in 5 μL/mg potassium phosphate buffer (pH 7), and then vortexed until homogenized. An equal volume of methanol containing 40 mM cysteine was added to promote conjugation of free, extracted isothiocyanate and incubated with rocking for 2 hours at room temperature. Samples were centrifuged at 14,000 *g* for 5 minutes, and then supernatant was filtered through a 0.45 μm PTFE filter for analysis by LC-MS/MS, as described below.

For analysis of serum samples, 50 μL of serum was mixed with 150 μL of acetonitrile to precipitate serum proteins. Samples were centrifuged at 10,000 *g* for 3 minutes, and then filtered through a 0.45 μm PTFE filter prior to analysis by LC-MS, as described below.

### LC-MS and LC-MS/MS analysis

#### Glucosinolate metabolism in spent bacterial media

Strain level variation in *Bt* ability to metabolize glucosinolate was analyzed by reversed phase liquid chromatography on an Agilent 1260 Infinity HPLC, using a 5 μm, 2 x 100 mm Gemini NX-C18 column (Phenomenex). For all samples, a volume of 5 μL was injected. Water with 0.1% (v/v) formic acid and acetonitrile with 0.1% (v/v) formic acid were used as mobile phase solvents, with a flow rate of 0.4 mL/min. Chromatographic separation was achieved using the following linear gradient (with percentages indicating levels of water with formic acid): 97% to 50%, 10 min; 50% to 5%, 1 min; 5%, 2 min; 5% to 97%, 1 min; 97%, 3 min. Coupled mass spectrometry data was collected with a Agilent 6520 quadrupole time-of-flight (qTOF) ESI mass spectrometer in positive polarity. Parameters for the 6520 qTOF MS were as follows: mass range, 50-1200 m/z; gas temperature, 350°C; drying gas flow rate, 11 L/min; nebulizer, 35 psig; fragmentor, 150 V; skimmer, 65 V. Glucosinolate conversion to ITC by *Bt* deletion mutants and complemented *Bf* strains, as well as *in vitro* protein activity, were analyzed by reversed phase liquid chromatography on an Agilent 1290 Infinity II HPLC, using a 1.8 μm, 2.1 x 50 mm Zorbax RRHD Eclipse Plus C18 column (Agilent). For all samples, a volume of 1 μL was injected. Water with 0.1% (v/v) formic acid and acetonitrile with 0.1% (v/v) formic acid were used as mobile phase solvents, with a flow rate of 0.6 mL/min. Chromatographic separation was achieved using the following linear gradient (with percentages indicating levels of water with formic acid): 95% to 5%, 4.2 min; 5% to 0%, 1 min; 0% to 95%, 0.4 min. Coupled mass spectrometry data was collected with either a Agilent 6545 quadrupole time-of-flight ESI mass spectrometer or an Agilent 6470 triple quadrupole (QQQ) mass spectrometer, indicated in figure legends as LC-MS or LC-MS/MS, respectively. For analyses with both instruments, data was collected in both positive and negative polarities. Parameters for the 6545 qTOF MS were as follows: mass range, 50-1700 m/z; gas temperature, 250C; drying gas flow rate, 12 L/min; nebulizer, 10 psig; fragmentor, 140 V; skimmer, 65 V. Parameters for the 6470 QQQ MS were as follows: gas temperature, 250C; gas flow rate, 12 L/min; nebulizer, 25 psig. Glucotropaeolin was detected using monitored transitions with the following parameters: polarity, negative; precursor ion, 408.04; product ions, 166 and 96.9; dwell, 100 ms; fragmentor, 138V; collision energy, 21V and 25V for respective product ions; cell accelerator, 4V. BITC-cys was detected using monitored transitions with the following parameters: polarity, positive; precursor ion, 271.1; product ions, 254 and 122; dwell, 100 ms; fragmentor, 100 V; collision energy, 6V and 10 V for respective product ions; cell accelerator, 4V.

#### Kinetic in vitro assays with purified protein

*In vitro* protein reactions were incubated at room temperature in the Agilent 1290 Infinity II autosampler and directly sampled with no quenching hourly by injection onto an Agilent 1290 Infinity II HPLC coupled to an Agilent 6545 qTOF MS. Injection volumes were 0.20 μL for all samples. No chromatography column was used. Water with 0.1% (v/v) formic acid and acetonitrile with 0.1% (v/v) formic acid were used as solvents, with the following isocratic flows (percentages indicate level of water with formic acid): 95% for 1 minute, 5% for 0.2 minutes. Mass spectrometry data was collected in fast polarity switching mode, with the following parameters: mass range, 50-700 m/z; drying gas temperature, 250C; drying gas flow rate, 12 L/min; nebulizer, 20 psig; fragmentor, 100V; skimmer, 50V.

#### Urine, feces, and serum sample analysis

Urine samples collected from mice following pure BGS doses were analyzed using the same method described above for glucosinolate metabolism in spent bacterial media, on the Agilent 6520 qTOF MS. Urine samples collected from mice following broccoli meals were analyzed by hydrophilic interaction liquid chromatography (HILIC) on an Agilent 1290 Infinity II HPLC, using a 1.7 μm, 2.1 x 50 mm Acquity BEH Amide column (Waters). For all samples, a volume of 1 μL was injected. Water with 0.125% (v/v) formic acid and 10 mM ammonium formate and acetonitrile were used as mobile phase solvents, with a flow rate of 0.6 mL/min. Chromatographic separation was achieved using the following linear gradient (with percentages indicating levels of water with formic acid and ammonium formate): 5%, 1.5 min; 5% to 62%, 4.5 min; 62%, 2 min; 62% to 71.5%, 0.5 min; 71.5% to 5%, 1 min, 5%, 2.5 min. Coupled mass spectrometry data was collected with an Agilent 6470 QQQ MS in both positive and negative polarities. Parameters for the 6470 QQQ MS were as follows: gas temperature, 250°C; gas flow rate, 12 L/min; nebulizer, 25 psig. Glucobrassicin was detected using monitored transitions with the following parameters: polarity, negative; precursor ion, 447.1; productions, 250.9 and 96.9; dwell, 50 ms; fragmentor, 156 V; collision energy, 22V and 26 V for respective product ions; cell accelerator, 4V. Glucoraphanin was detected using monitored transitions with the following parameters: polarity, negative; precursor ion, 436.0; product ions, 372 and 96.9; dwell, 50 ms; fragmentor, 152 V; collision energy, 18V and 26 V for respective product ions; cell accelerator, 4V. Sulforaphane N-acetyl cysteine conjugate (SFN-NAC) was detected using monitored transitions with the following parameters: polarity, positive; precursor ion, 341.1; product ions, 177.9 and 114; dwell, 50 ms; fragmentor, 160V; collision energy, 10 V and 22 V for respective product ions; cell accelerator, 4V. Sulforaphane cysteine conjugate (SFN-cys) was detected using monitored transitions with the following parameters: polarity, positive; precursor ion, 299.1; product ions, 136 and 114; dwell, 50 ms; fragmentor, 100V; collision energy, 6 V and 22 V for respective product ions; cell accelerator, 4V. Benzyl isothiocyanate N-acetyl cysteine conjugate (BITC-NAC) was detected using monitored transitions with the following parameters: polarity, positive; precursor ion, 313.1; product ions, 164 and 91.1; dwell, 50 ms; fragmentor, 100V; collision energy, 6 V and 42 V for respective product ions; cell accelerator, 4V. Glucotropaeolin and BITC-cys were detected using monitored transitions as described above for detection in spent bacterial media.

Fecal samples collected from mice following both pure glucotropaeolin doses and broccoli meals were analyzed by reversed phase chromatography, using the same chromatography column and method used to determine glucosinolate metabolism in spent bacterial media. Coupled mass spectrometry data was collected with an Agilent 6470 QQQ MS in both positive and negative polarities, with the same parameters as described above. BITC-cys and BGS were detected using monitored transitions as described above for detection in spent bacterial media. Following analysis, fecal samples resulting from broccoli meals were diluted 10 times in HILIC acetonitrile mobile phase, filtered through a 0.45μm PTFE filter, and further analyzed using HILIC, using the same chromatography column and method used to separate metabolites in urine samples resulting from broccoli meals. Coupled MS data was collected with an Agilent 6470 QQQ MS in both positive and negative polarities, with the same parameters as described above. BITC-cys, SFN-cys, glucotropaeolin, and glucoraphanin were detected using monitored transitions as described above. Mouse serum samples collected post-sacrifice were analyzed using HILIC, with the same chromatography column and method used to separate metabolites in urine samples resulting from broccoli meals. Coupled MS data was collected using an Agilent 6545 qTOF MS in both positive and negative polarities, with the same parameters as described above.

#### Data analysis

Extracted ion chromatograms (EIC) were extracted from raw data files using MassHunter Qualitative Data Analysis software. For MS data collected using the Agilent 6545 qTOF, EICs were extracted using a 40 ppm window centered on the exact m/z value. For data collected using the Agilent 6520 qTOF, a 20 ppm window was used. Exact m/z values for the different substrates and products are as follows, with (+) or (−) indicating detection of the [M+H]^+^ or [M-H]^−^ species in positive or negative polarity, respectively: glucotropaeolin, (−) 408.0428; glucoraphanin, (−) 436.0411; cellobiose and maltose, (−) 341.1089; glucobrassicin, (−) 447.0538; hexose, (−) 179.0561; sulforaphane, (+) 178.0355; BITC-cys, (+) 271.0570; BITC-NAC, (+) 313.0675; SFN-cys, (+) 299.0552; SFN-NAC, (−) 341.0658.

